# DeepSTARR predicts enhancer activity from DNA sequence and enables the *de novo* design of enhancers

**DOI:** 10.1101/2021.10.05.463203

**Authors:** Bernardo P. de Almeida, Franziska Reiter, Michaela Pagani, Alexander Stark

## Abstract

Enhancer sequences control gene expression and comprise binding sites (motifs) for different transcription factors (TFs). Despite extensive genetic and computational studies, the relationship between DNA sequence and regulatory activity is poorly understood and enhancer *de novo* design is considered impossible. Here we built a deep learning model, DeepSTARR, to quantitatively predict the activities of thousands of developmental and housekeeping enhancers directly from DNA sequence in *Drosophila melanogaster* S2 cells. The model learned relevant TF motifs and higher-order syntax rules, including functionally non-equivalent instances of the same TF motif that are determined by motif-flanking sequence and inter-motif distances. We validated these rules experimentally and demonstrated their conservation in human by testing more than 40,000 wildtype and mutant *Drosophila* and human enhancers. Finally, we designed and functionally validated synthetic enhancers with desired activities *de novo*.

---

Enhancers^1^ are genomic elements that regulate the cell type-specific transcription of target genes, thereby controlling animal development and physiology^2^. A characteristic feature of enhancers is their ability to activate transcription outside their endogenous genomic contexts^3^, which suggests that all the necessary cis-regulatory information is contained within the enhancers’ DNA sequences. Indeed, enhancer sequence mutations can drastically alter enhancer function and are associated with developmental defects^2^, morphological evolution^4^, and human disease^5^.

Enhancers typically contain multiple sequence motifs that are binding sites for sequence-specific transcription factors (TFs)^6^. Understanding how motifs and their arrangements (i.e. their number, order, orientation and spacing – termed here collectively *motif syntax*) relate to enhancer function has remained one of the most important open questions in modern biology. Systematic mutagenesis of various individual enhancers revealed a complex picture, whereby changing nucleotides or altering motif syntax affected the function of some enhancers but not others^7–26^. These contradictory observations made it difficult to define the relationships between enhancer sequence and function^18, 27^.

Many computational approaches have sought to predict enhancer activities from DNA sequences using local DNA features, e.g. motif dictionaries or *de-novo k-*mers, and selected syntax rules in various thermodynamic or machine-learning frameworks^16, 17, 26, 28–39^. Despite remarkable success, these approaches did not reveal how the elements of motif syntax collaborate to determine enhancer activity. In addition, they did not consider the mutual compatibilities between certain enhancer- and promoter types recently reported for different transcriptional programs^40–42^. Thus, quantitatively predicting the regulatory activity of enhancers and the *de novo* design of synthetic enhancers have remained open challenges for decades.

Previous approaches typically modelled enhancer sequences explicitly via pre- defined sets of features, which were informed by prior biological knowledge^43^. In contrast, deep learning, in particular convolutional neural networks, do not require prior knowledge and can learn accurate models directly from raw data^44–53^. Once trained on raw data, these models allow the extraction and interpretation of the learned rules by novel types of tools^44, 45, 47, 48, 54–60^. For example, when applied to ChIP-nexus data that measures TF-binding genome-wide at high resolution, a convolutional neural network was able to learn motifs and syntax rules for cooperative TF binding^47^. Similarly, this approach was used to model DNA accessibility^45, 46, 52, 59^, transcriptional reporter activities^51^ and predict genetic variant effects^53^. Nevertheless, an ultimate sequence-to-enhancer activity model that learns the cis-regulatory code to quantitatively predict enhancer activities in a single cell type is still missing.

Here, we built a new deep learning model, DeepSTARR, to predict enhancer activity towards two promoters from the distinct developmental and housekeeping transcriptional programs in *Drosophila melanogaster* S2 cells directly from the DNA sequence. For both programs, DeepSTARR quantitatively predicts enhancer activity for unseen sequences and reveals different coding features for the two programs, including specific TF motifs that we validate experimentally. We further extract motif syntax rules, including favorable and unfavorable sequence contexts and inter-motif distances, which are predictive of enhancer activity in *Drosophila* and human enhancers, as we validate experimentally by high-throughput mutagenesis of thousands of enhancers and enhancer variants. These rules allowed the design of synthetic enhancers with desired activity levels *de novo*.

## Results

### DeepSTARR quantitatively predicts enhancer activity from DNA sequence

To learn the cis-regulatory information encoded in enhancer sequences in an unbiased way, we developed a new deep learning model called DeepSTARR that predicts enhancer activity directly from DNA sequence. First, we used UMI-STARR-seq^61, 62^ to generate genome-wide high resolution, quantitative activity maps of developmental and housekeeping enhancers, representing the two main transcriptional programs in *Drosophila* S2 cells^40–42^ (Fig 1A). We identified 11,658 developmental and 7,062 housekeeping enhancers (Fig 1B, S1A,B). These enhancers are largely non-overlapping, confirming the specificity of the different transcriptional programs^40–42^. These genome- wide enhancer activity maps provide a high-quality dataset to build predictive models of enhancer activity and characterize the sequence determinants of two major enhancer types.

**Figure 1.**
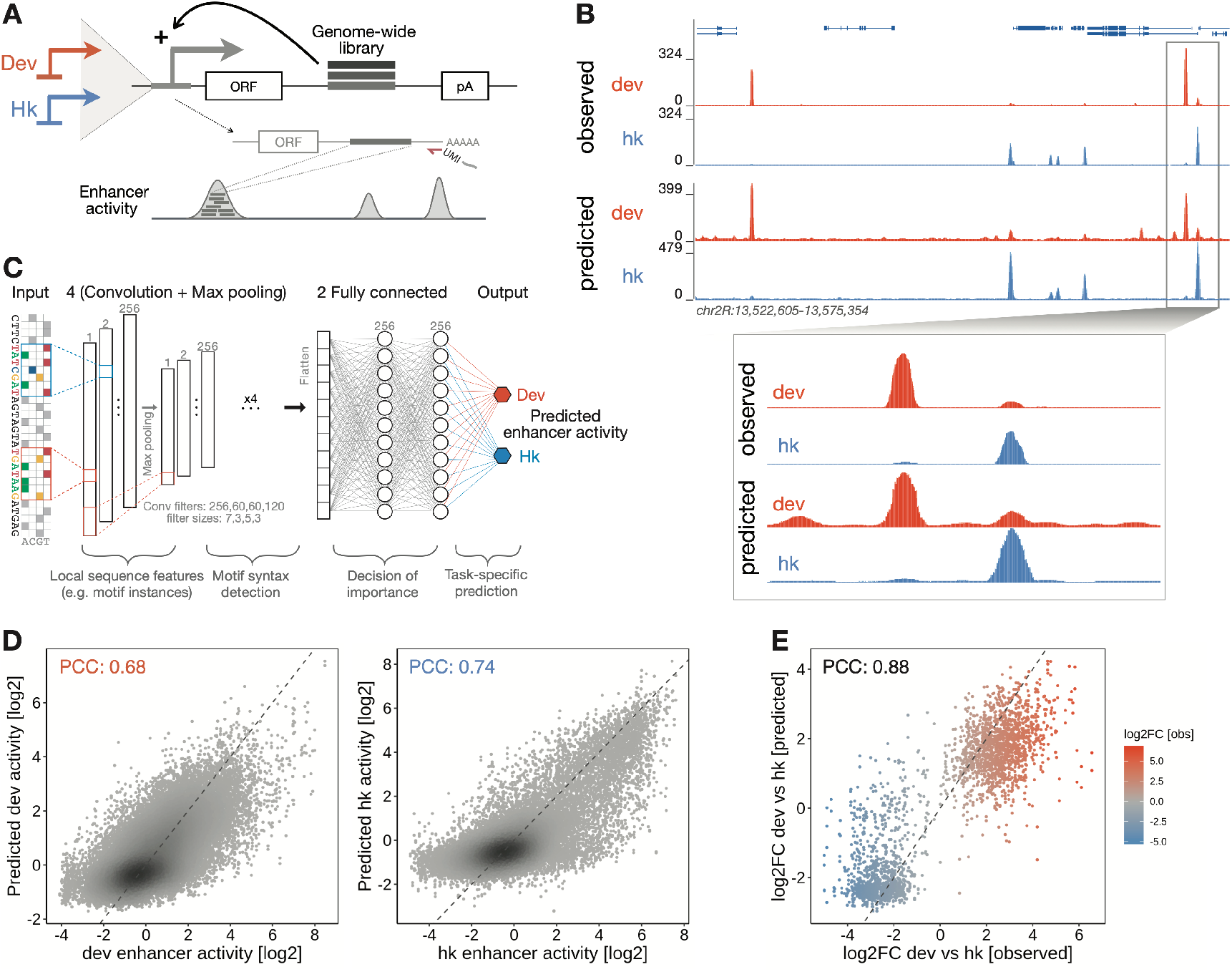
DeepSTARR quantitatively predicts enhancer activity genome-wide from DNA sequence. **A)** Schematics of genome-wide UMI-STARR-seq using the developmental (Drosophila synthetic core promoter (DSCP); red) and housekeeping (RpS12; blue) promoters. **B)** DeepSTARR predicts enhancer activity genome-wide. Genome browser screenshot depicting UMI-STARR-seq observed and predicted profiles for both promoters for a locus on held-out test chromosome 2R. **C)** Architecture of the multi-task convolutional neural network DeepSTARR that was trained to simultaneously predict quantitative developmental and housekeeping enhancer activities (UMI-STARR-seq) from 249 bp DNA sequences. **D)** DeepSTARR predicts enhancer activity quantitatively. Scatter plots of predicted vs. observed developmental (left) and housekeeping (right) enhancer activity signal across all DNA sequences in the test set chromosome. Color reflects point density. **E)** DeepSTARR quantitatively predicts developmental and housekeeping enhancer–promoter specificity. Predicted vs. observed log2 fold-change (log2FC) between developmental and housekeeping activity for all enhancer sequences in the test set chromosome. PCC: Pearson correlation coefficient.

We built the multi-task convolutional neural network DeepSTARR to map 249 bp long DNA sequences tiled across the genome to both their developmental and their housekeeping enhancer activities (Fig 1C). We adapted the Basset convolutional neural network architecture^45^ and designed DeepSTARR with four convolution layers, each followed by a max-pooling layer, and two fully connected layers (Fig 1C). The convolution layers identify local sequence features (e.g. TF motifs) and increasingly complex patterns (e.g. TF motif syntax), while the fully connected layers combine these features and patterns to predict enhancer activity separately for each enhancer type.

We evaluated the predictive performance of DeepSTARR on a held-out test chromosome. The predicted and observed enhancer-activity profiles were highly similar for both developmental (Pearson correlation coefficient (PCC)=0.68) and housekeeping (PCC=0.74) enhancers (Fig 1B,D, S1). This performance is close to the concordance between experimental replicates (PCC=0.73 and 0.76, respectively; Fig S1C), suggesting that the model accurately captures the regulatory information present in the sequences and the differences between developmental and housekeeping enhancers (Fig 1E). DeepSTARR performed better than methods based on known TF motifs or unbiased *k*- mer counts^39^, both at predicting continuous enhancer activity and at binary classification of enhancer sequences (Fig S1D). Thus, DeepSTARR learned generalizable features and rules *de novo* directly from the DNA sequence that allow the prediction of enhancer activities for unseen sequences.

### DeepSTARR reveals important TF motif types required for enhancer activity

In order to understand the features and rules learned by DeepSTARR, we quantified how each individual nucleotide in every sequence contributes to the predicted developmental and housekeeping enhancer activities^47, 55, 63, 64^ (Fig 2A; see Methods), and consolidated recurrent highly scoring sequence patterns into motifs ^56^. This uncovered distinct TF motifs that are known to occur in developmental and housekeeping enhancers^26, 40^, thus validating the approach and reinforcing the mutual incompatibility of the two transcriptional programs (Fig 2A,B, S2).

**Figure 2.**
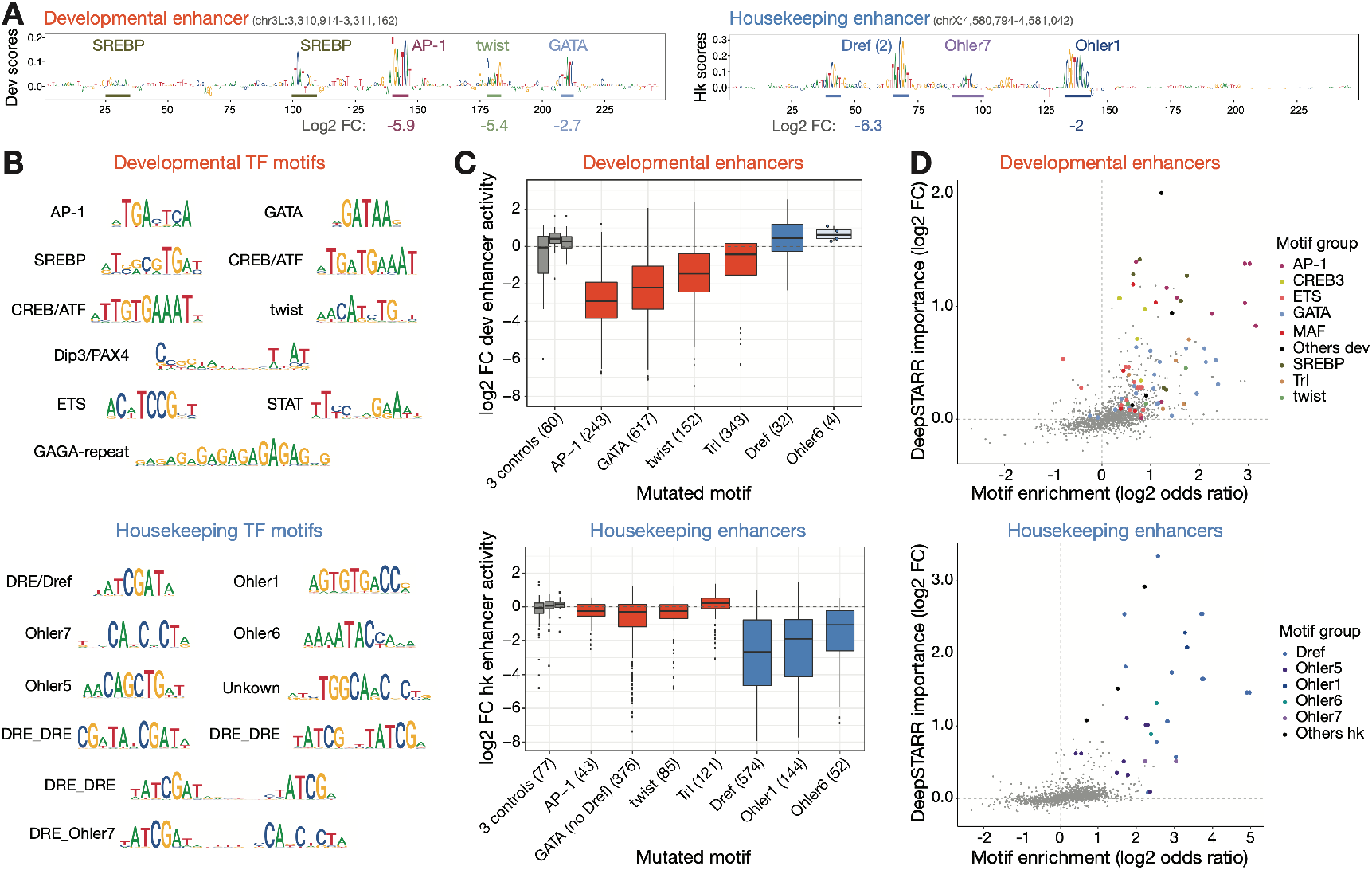
DeepSTARR reveals important TF motif types that we validate experimentally. **A)** DeepSTARR derived developmental and housekeeping nucleotide contribution scores for strong developmental (left) and housekeeping (right) enhancer sequences, respectively. Regions with high scores resembling known TF motifs are highlighted. Log2 fold-change values (log2FC; bottom) indicate the impact on enhancer activity of mutating all instances of each motif type. **B)** DeepSTARR motifs discovered by TF–Modisco by summarizing recurring predictive sequence patterns from the sequences of all developmental (top) and housekeeping (bottom) enhancers and their associated nucleotide contribution scores. **C)** Developmental and housekeeping TF motifs are specifically required for the respective enhancer types. Enhancer activity changes (log2 FC) for developmental (top) and housekeeping (bottom) enhancers after mutating all instances of three control motifs (grey), four predicted developmental motifs (AP-1, GATA, twist, Trl; red) and three predicted housekeeping motifs (Dref, Ohler1, Ohler6; blue). Number of enhancers mutated for each motif type are shown. The box plots mark the median, upper and lower quartiles and 1.5× interquartile range (whiskers); outliers are shown individually. **D)** DeepSTARR discovers important TF motifs not obvious by motif enrichment. Comparison between motif enrichment (log2 odds ratio; x-axis) and DeepSTARR’s predicted global importance (y-axis) for all representative TF motifs (Fig S2) in developmental (top) and housekeeping (bottom) enhancers. Important motifs for each enhancer type are highlighted.

We experimentally tested the requirements of select TF motifs for enhancer activity across hundreds of enhancers by performing large-scale motif mutagenesis (3,415 motif- mutations in 804 developmental and 872 housekeeping enhancers; Fig. 2C, S3). Consistent with their predicted importance, mutating four developmental motifs (GATA, AP-1, twist, Trl) substantially reduced the activity of developmental but not housekeeping enhancers, with AP-1 and GATA motifs being the most important, as predicted by DeepSTARR. In contrast, mutating three housekeeping motifs (Dref, Ohler1, Ohler6) affected only housekeeping enhancers, and mutating three control motifs (length-matched random motifs to control for enhancer-sequence perturbation) did not have any impact. For example, GATA motifs were only important when present in developmental but not housekeeping enhancers, whereas the opposite was true for Dref motifs (Fig. 2C). Interestingly, the motifs learned by DeepSTARR were not restricted to highly abundant motifs but included other motifs such as SREBP, CREB and ETS motifs, which were not or only weakly enriched in S2 developmental enhancers and could not have been found by methods based on over-representation (Fig. 2B,D). Even for more abundant motifs, motif enrichment did not always predict motif importance (Fig. 2D), i.e. the DeepSTARR score of the motif embedded in 100 random DNA sequences (see Methods and ref. ^58^). This shows that DeepSTARR can discover motifs, and likely other sequence features, that are relatively rare in enhancers but still important for enhancer activity.

### Non-equivalent instances of the same TF motif

Since enhancers often contain multiple instances of the same motif type, we next assessed the contribution of each individual instance of the GATA, AP-1, twist, Trl, and Dref motifs by DeepSTARR (Fig S5A) and by experimental mutagenesis (Fig S3A, S5B). Unexpectedly, individual instances of the same motif were frequently predicted and experimentally validated to have distinct contributions to enhancer activities, both across different enhancers and within the same enhancer (Fig 3A-C, S5).

**Figure 3.**
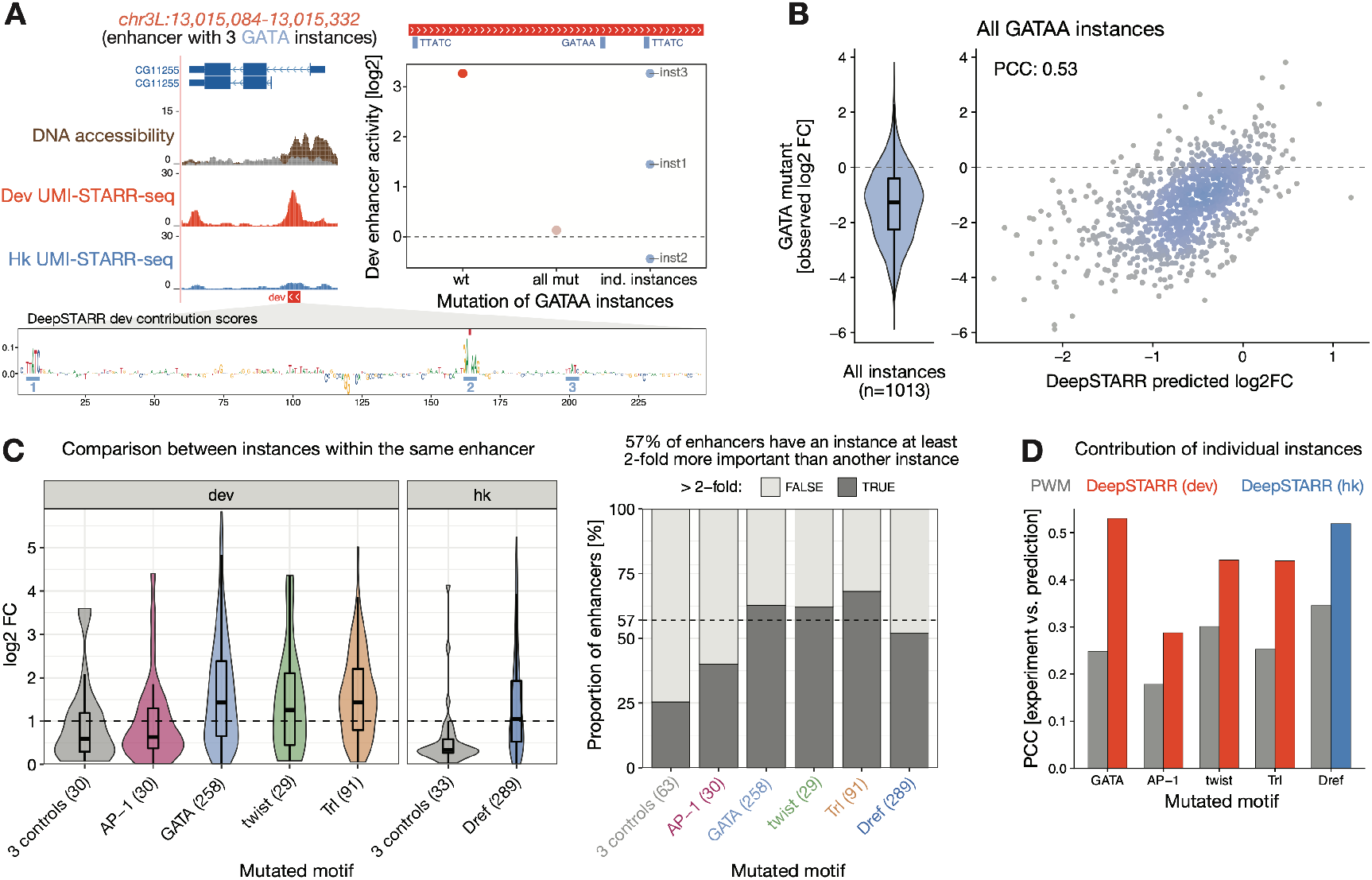
Instances of the same TF motif have non-equivalent contributions to enhancer activity. **A)** Developmental enhancer with three non-equivalent GATA instances. Left: Genome browser screenshot showing tracks for DNA accessibility (from ^61^) and developmental and housekeeping UMI-STARR-seq for the *CG11255* locus. The designed oligo covering the enhancer selected for motif mutagenesis is shown. Right: log2 activity of the wildtype enhancer compared with the activity when all GATA instances are simultaneously mutated or each individual instance at a time. Bottom: DeepSTARR nucleotide contribution scores for the same developmental enhancer with the three GATA instances highlighted. **B)** DeepSTARR predicts the contribution of individual GATA instances. Distribution of experimentally measured enhancer activity fold-change (log2 FC) after mutating 1,013 different GATA instances across developmental enhancers (violin plot), compared with the log2 FC predicted by DeepSTARR. The box plots mark the median, upper and lower quartiles and 1.5× interquartile range (whiskers). **C)** Different instances of the same TF motif in the same enhancer are not equivalent. Left: Distribution of enhancer activity change (log2 FC) between mutating the least and the most important instance of each motif type per enhancer. Dashed line represents 2-fold difference between instances in the same enhancer. Right: Proportion of enhancers with two or more instances that have an instance at least 2-fold more important than another instance (dark grey). Dashed line represents the average across the different motif types (excluding control motifs): 57% of enhancers. Number of enhancers mutated for each motif type are shown. Box plots as in (B). **D)** DeepSTARR predicts motif instance contribution better than position weight matrix (PWM) motif scores. Bar plots showing the PCC between predicted (by DeepSTARR or PWM) and observed log2 fold-change for mutating individual instances of each motif type.

The enhancer shown in Fig 3A for example contains three GATA instances with very different contributions as predicted and determined experimentally: the second instance is the most important one, followed by the first, and the third. The agreement between predictions and experiments holds across all 1,013 GATA instances tested (PCC=0.53; Fig 3B) and the non-equivalency of motif instances is widespread: 57% of enhancers with multiple instances had motifs with >2-fold and 70% with >1.5-fold differences (Fig 3C). These differences are not well captured by existing position weight matrix (PWM) motif scores (Fig 3D, S6), suggesting that the importance of motif instances depend on complex sequence features outside the core motif. The observation that different instances of the same motif type (with identical sequences) can have vastly different contributions to enhancer activity is an important underappreciated phenomenon that complicates our understanding of enhancer sequences and non-coding variants (see Discussion).

### Flanking sequence influences the importance of TF motifs

To explore the syntax features that affect the importance of a motif instance, we examined the motif flanking nucleotides, which can contribute to enhancer activity^12, 13, 18, 29, 65–69^. For each motif type, we sorted all instances by their predicted importance to determine the optimal flank length and sequence (Fig 4A,B, S7A). For example, important GATAA sequences had a G at position +1, whereas non-important ones had a T at position +1 and a G at position -1 (Fig 4A). In contrast, up to 5 bp flanking up- and down-stream affected the importance of Trl instances, with flanking GA-repeats correlating with increased importance (Fig 4A). The flanks of high and low importance motif instances predicted by DeepSTARR were largely concordant with those identified by motif mutagenesis (Fig 4B, S7A) and refine known PWM models for the predicted TFs (Fig 4B).

**Figure 4.**
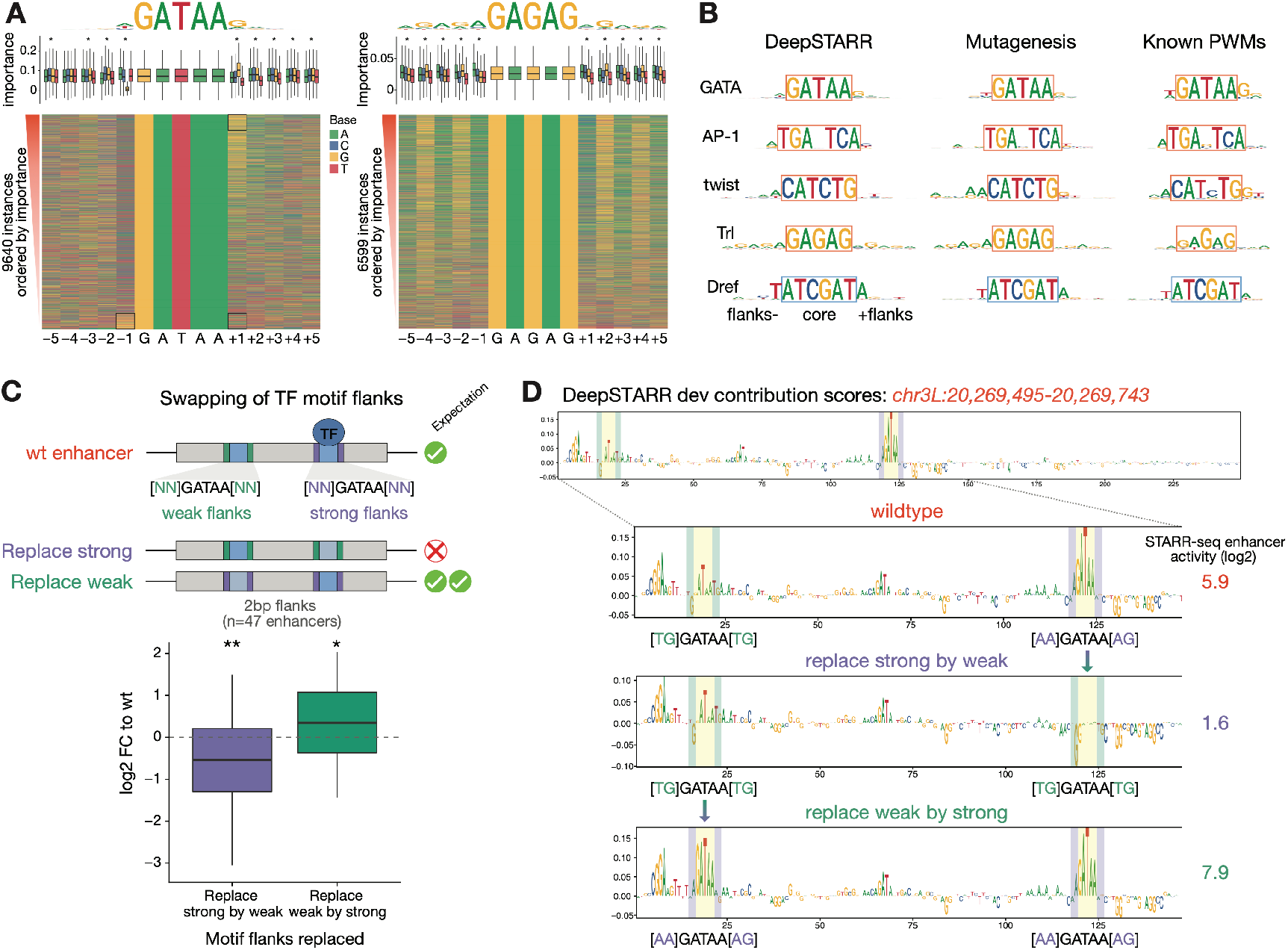
Contribution of TF motifs depends on the flanking sequence. **A)** Motif contribution correlates with flanking base-pairs. Heatmap: Flanking nucleotides of GATAA (GATA; left) and GAGAG (Trl; right) instances across developmental enhancers sorted by their DeepSTARR predicted contribution. Box plots: Importance of motif instances according to the different bases at each flanking position. * marks positions with significant differences between the four nucleotides (FDR-corrected Welch One-Way ANOVA test p-value < 0.01). The box plots mark the median, upper and lower quartiles and 1.5× interquartile range (whiskers). Top: logos of the top 90^th^ percentile motif instances. **B)** Length and identity of flanks differ between motif types. Comparison of optimal motif logos (top 90^th^ percentile motif instances) as predicted by DeepSTARR or measured experimentally by motif mutation, with the PWM logos existing in *Drosophila* TF databases. Note that DeepSTARR and mutagenesis motif instances were selected to all contain the same core sequence and therefore only differ in their flanking sequence. **C)** GATA flanking nucleotides are sufficient to switch motif contribution in 47 developmental enhancers that contain one strong (purple) and one weak (green) GATA instance (≥ 2-fold difference between instances as assessed by mutagenesis). Enhancer activity change (log2 FC) when 2 bp flanks of strong instances were replaced by the flanks of weak instances (purple) and vice versa (green). ** p-value < 0.01, * < 0.05 (Wilcoxon signed rank test). Box plots as in (A). **D)** Example of a developmental enhancer with one weak (TGGATAATG; green) and one strong (AAGATAAAG; purple) GATA instance. DeepSTARR nucleotide contribution scores and UMI-STARR-seq measured enhancer activity (log2; on the right) are shown for the wildtype sequence (top) and for the sequences where the 2 bp flanks of the strong instance were replaced by the ones of the weak instance (middle) and vice versa (bottom).

To experimentally validate the functional contribution of motif flanking sequence predicted by DeepSTARR, we swapped the flanking nucleotides of strong and weak GATA instances (≥ 2-fold difference as assessed by mutagenesis) in 47 enhancers (Fig 4C). Indeed, replacing the 2 bp flanks of strong instances by the flanks of weak instances reduced enhancer activity, whereas replacing the flanks of weak instances by the flanks of strong ones increased enhancer activity (Fig 4C, S7B). DeepSTARR recapitulated the observed effects, i.e. the addition of weak flanks converted a strong GATA instance to a weak one as indicated by the decreased contribution at the nucleotide level, and vice versa for a weak instance that was converted to a strong one (Fig 4D). Swapping 5 bp flanks yielded consistent results with slightly stronger effects (Fig S7B). In addition, swapping the flanks was sufficient to switch motif contributions, as determined by subsequent motif mutagenesis (Fig. S7B)). Thus, as DeepSTARR is not biased by prior knowledge about TF motifs but is trained on DNA sequence alone, it can not only identify important motif types but also refine the optimal flanking sequence. Experimentally, we confirm that the flanking sequence can be sufficient to switch motif contribution and should be considered when assessing motif importance or the impact of motif-disrupting mutations.

### *In silico* analysis reveals distinct modes of motif cooperativity

The distance between TF motifs is thought to be important for TF cooperativity^6, 13, 18, 47, 70–73^. To determine how the distance between motifs contributes to enhancer activity, we interrogated DeepSTARR to uncover potential preferences in TF motif distance in enhancer sequences. We analyzed *in silico* how predicted enhancer activity is affected by the relative distance between two motif instances (*MotifA*/*MotifB*), following a strategy adapted from^47^ (Fig 5A, S8A): we embedded *MotifA* in the center of synthetic random DNA sequences and *MotifB* at a range of distances from *MotifA*, both up- and downstream. We then predicted the activity of the different synthetic sequences using DeepSTARR and calculated a cooperativity score for each motif pair, where a value higher than 1 means positive synergy (Fig 5A, S8A).

**Figure 5.**
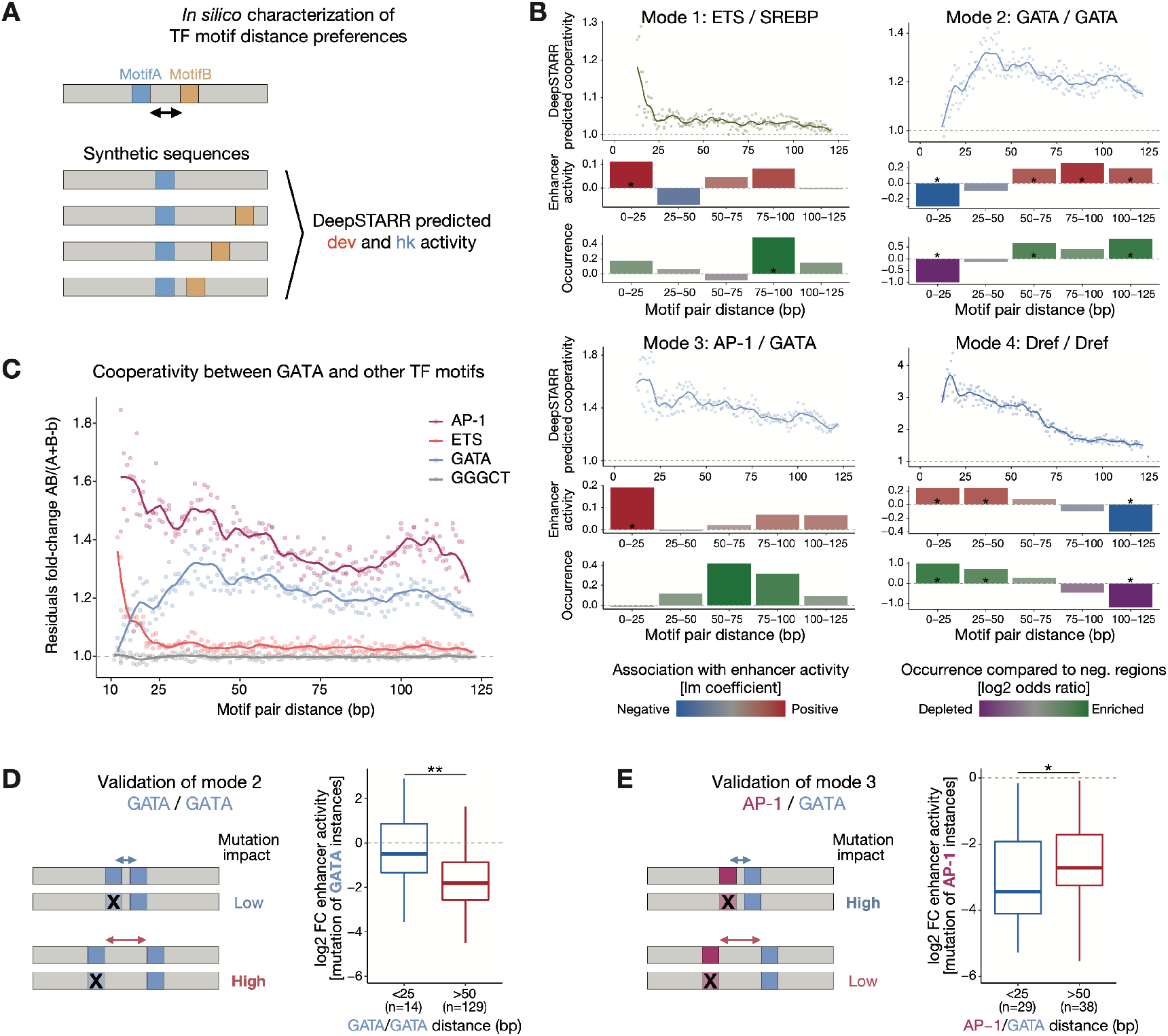
*In silico* analysis reveals distinct modes of motif cooperativity. **A)** Schematic of *in silico* characterization of TF motif distance preferences. *MotifA* was embedded in the center of 60 synthetic random DNA sequences and *MotifB* at a range of distances from *MotifA*, both up- and downstream. Both the average developmental and housekeeping enhancer activity is predicted by DeepSTARR. The cooperativity (residuals) between *MotifA* and *MotifB* as a function of distance is quantified as the activity of *MotifA+B* divided by the sum of the marginal effects of *MotifA* and *MotifB* (*MotifA* + *MotifB –* backbone (b)) (see Methods). **B)** DeepSTARR predicts distinct modes of motif cooperativity. Show for example motif pairs: ETS/SREBP (mode 1), GATA/GATA (2), AP-1/GATA (3) and Dref/Dref (4). Top: Cooperativity between two motif instances at different distances. Points showing the median interaction across all 60 backbones for each motif pair distance (both up- and downstream distances are combined) together with smooth lines; dashed line at 1 represents no synergy. Middle: Association between enhancer activity and the distance at which the motif pair is found. Coefficient (y-axis) and p-value from a multiple linear regression including, as independent variables, the number of instances for the different developmental or housekeeping TF motif types. Bottom: Motifs are often at suboptimal distances in developmental enhancers. Odds ratio (log2) by which the two motifs are found within a specified distance from each other in enhancers compared with negative genomic regions. Color legend is shown. * FDR-corrected Fisher’s Exact test p-value < 0.05. **C)** Cooperativity between three motif types (and GGGCT as control) with a central GATA motif in developmental enhancers at different distances to the GATA motif. Points showing the median interaction across all 60 backbones for each motif pair distance (both up- and downstream distances are combined) together with smooth lines; dashed line at 1 represents no synergy. **D)** Motif mutagenesis validates that GATA instances distal to a second GATA instance are more important. Left: expected mutational impact when mutating GATA instances depending on the distance to other GATA motifs. Right: enhancer activity changes (log2 FC) after mutating GATA instances at suboptimal close (< 25 bp) or optimal longer (> 50 bp) distance to a second instance. Number of instances are shown. ** p- value < 0.01 (Wilcoxon rank-sum test). The box plots mark the median, upper and lower quartiles and 1.5× interquartile range (whiskers). **E)** Motif mutagenesis validates that AP-1 instances closer to a second GATA instance are more important. Same as in (D). * p-value < 0.05 (Wilcoxon rank-sum test).

Motif distances had indeed a strong influence on predicted enhancer activity and we observed four distinct modes of distance-dependent TF motif cooperativity: motif pairs can synergize exclusively at close distances (< 25 bp; mode 1), exclusively at longer distances (> 25 bp; 2), preferentially at closer distances and either plateau (3) or decay (4) at long distances (> 75 bp; Fig 5B, S8B-D). While all motifs in housekeeping enhancers cooperate according to mode 4 (decay), modes 1 to 3 all occur for motifs in developmental enhancers (Fig S8C,D). Interestingly, whether cooperativity followed modes 1, 2 or 3 depended on the motif pair and even changed for a given motif based on the partner motif (Fig 5C, S8C). For example, GATA/ETS synergized only when closer than 25 bp (mode 1), whereas GATA/GATA synergy was lost at short distances (mode 2) and GATA/AP-1 cooperated according to mode 3 (Fig 5C). Thus, DeepSTARR predicts distinct modes of motif cooperativity that can determine the contribution of different motif instances.

We next asked how frequently these optimal inter-motif distances occur in endogenous enhancers compared to negative regions. Motif pairs of housekeeping enhancers followed the optimal spacing rules (enrichment at close distances; Fig 5B, S9A,D), as did some motif pairs in developmental enhancers such as GATA/GATA motif pairs that were strongly depleted at close and enriched at longer distances (Fig. 5B).

However, several pairs in developmental enhancers occurred only rarely at optimal distances (e.g. ETS/SREBP and AP-1/GATA; Fig 5B, S9A,C), even though the enhancer activities followed the predicted optimal spacing rules also in these cases (Fig 5B, S9). For instance, even though ETS/SREBP motifs separated by short distances (< 25 bp) were rare, such motif pairs were associated with stronger enhancer activity than pairs separated by larger distances (75-100 bp; Fig 5B), validating the ETS/SREBP motifs’ optimal distance.

To experimentally test the importance of motif pairs at optimal versus non-optimal distances more directly, we mutated either GATA or AP-1 motifs at close (< 25 bp) and longer distances (> 50 bp) to a GATA instance (Fig 5D,E). The results validated the DeepSTARR predictions and showed higher importance of GATA/GATA pairs at longer (Fig 5D) and AP-1/GATA pairs at closer distances (Fig 5E). Thus, different motif pairs display distinct distance preferences, which dictate the contribution of individual motif instances to overall enhancer activity. As endogenous enhancers often contain motif pairs at non-optimal distances, optimal distances only become apparent by our *in silico* analysis but not in frequency-based analyses.

### Motif syntax rules are generalizable to human enhancers

To test if individual instances of the same motif also contribute differently to enhancer activities in humans and if motif flanks and spacing determine the different contributions, we chose the human colon cancer cell line HCT116 as a model. We selected nine TF motifs based on motif enrichment analysis (AP-1, P53, MAF, CREB1, ETS, EGR1, MECP2, E2F1 and Ebox/MYC), mutated all their instances in 1,083 enhancers and assessed the enhancer activity of wildtype and mutant sequences by UMI-STARR-seq (Fig S10; see Methods). This revealed that AP-1 and P53 motifs were the most important motifs (median 5.6- and 5.5-fold reduction, respectively), followed by MAF (3.1), CREB1 (2), ETS (1.9) and EGR1 (1.5), while MeCP2, E2F1 and Ebox/MYC motifs had the least impact on enhancer activity (lower than 1.5-fold; Fig S10D-F). Based on these results, we chose AP-1, P53, MAF, CREB1, ETS and EGR1 motifs for the analysis of motif instances.

Mutation of hundreds of individual motif instances showed that instances of the same TF motif are not functionally equivalent (Fig 6A-C, S11A). For example, the enhancer shown in Fig 6A contains four AP-1 instances with very different contributions to enhancer activity as judged by fold-changes after motif instance mutagenesis between 1.2- and 3.8-fold. Interestingly, DNase I footprinting data from a related colon cancer cell line (RKO^76^) suggests that the AP-1 instance with low importance was not bound endogenously, in contrast to the three important AP-1 instances (Fig 6A). Both results generalize to all tested motifs and across enhancers: 57% of human enhancers displayed non-equivalent instances of the same motif type (Fig 6B,C) and TF motif instances with DNase I footprints are more important than those without (Fig 6D), supporting the functional differences between motif instances at endogenous enhancers.

**Figure 6.**
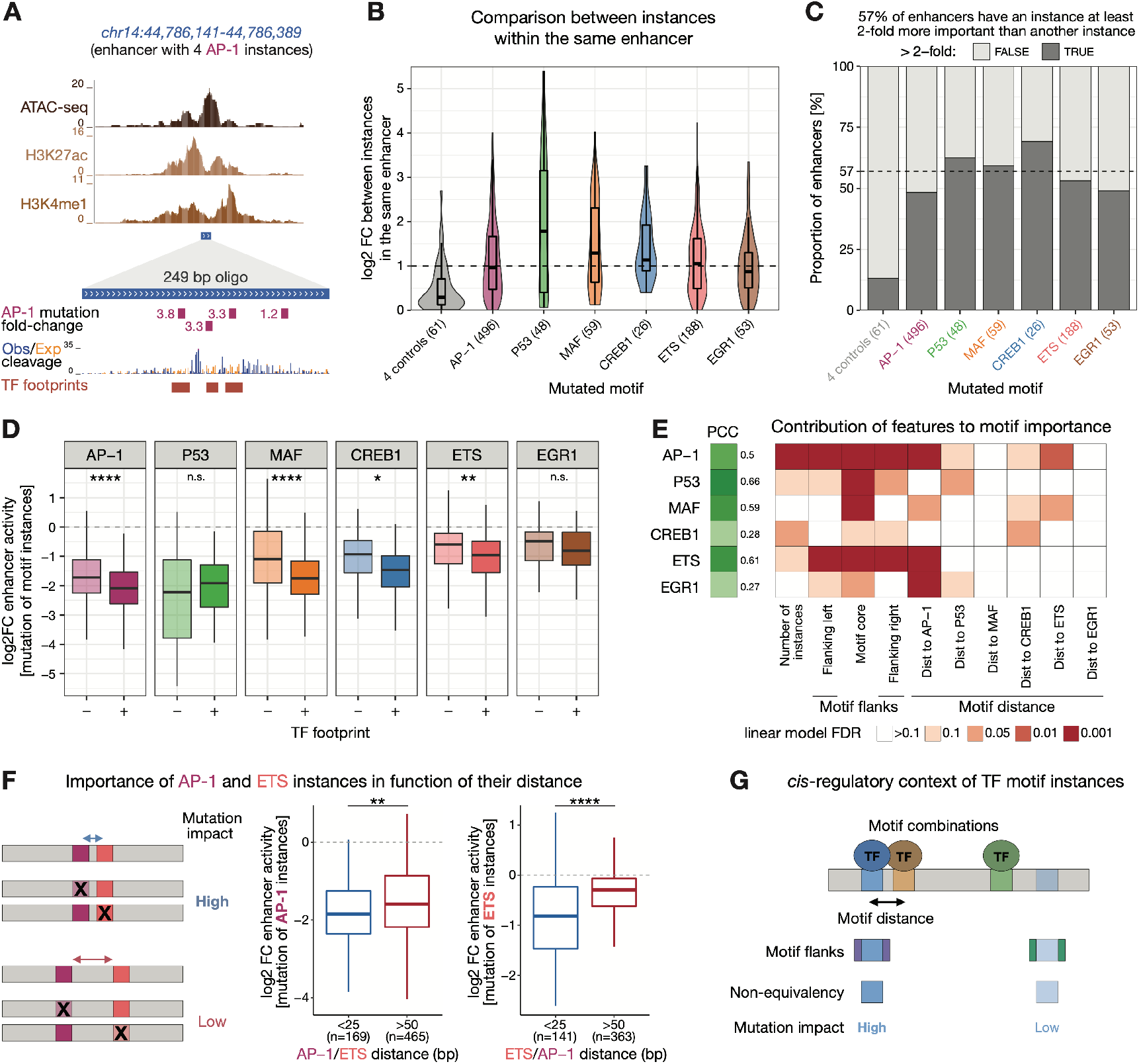
Motif syntax rules dictate the contribution of TF motif instances in human enhancers. **A)** Top: Genome browser screenshot showing DNA accessibility (ATAC-seq data from^74^) and enhancer chromatin marks (H3K27ac and H3K4me1 from ENCODE^75^) for a human HCT116 enhancer (chr14:44,786,141-44,786,389) with 4 AP-1 instances. Bottom: The designed 249 bp oligo covering the enhancer summit used for motif mutagenesis is shown together with its containing AP-1 motif instances and the impact on enhancer activity (negative fold-change) of mutating each individual instance. Observed and expected per-nucleotide DNase I cleavage and consensus TF footprints from a related colon cancer cell line (RKO; data from ^76^) are shown below. **B)** TF motif non-equivalence is widespread in human enhancers. Distribution of log2 FC enhancer activity between mutating the least and the most important instance of each motif type per enhancer. Dashed line represents 2-fold difference between instances in the same enhancer. Number of enhancers mutated for each motif type are shown. The box plots mark the median, upper and lower quartiles and 1.5× interquartile range (whiskers). **C)** 57% of enhancers have a motif instance that is at least 2-fold more important than another instance. Grey bars: proportion per motif type; dashed line: average across motif types (excluding control motifs). **D)** Important TF motif instances are associated with TF footprints. Log2 FC enhancer activity of mutating individual instances that do not (-) or do (+) overlap TF footprints (FDR 0.001) in RKO cells (DNase I footprinting data from^76^). **** p-value < 0.0001, ** < 0.01, * < 0.05, n.s. non-significant (Wilcoxon rank-sum test). Box plots as in (B). **E)** Motif syntax rules dictate the contribution of TF motif instances in human enhancers. For each TF motif type (rows), we built a linear model containing the number of instances, the motif core (defined as the nucleotides included in each TF motif PWM model) and flanking nucleotides, and the distance to all other TF motifs to predict the contribution of its individual instances (mutation log2 fold-change, from Fig S11A) across all enhancers. The PCC between predicted and observed motif contribution is shown with the green color scale on the left. Heatmap shows the contribution of each feature (columns) for each model, colored by the FDR-corrected p-value (red scale). **F)** Motif mutagenesis shows that AP-1 and ETS instances closer to each other are more important to enhancer activity. Left: expected mutational impact when mutating AP-1 and ETS instances depending on the distance to each other. Middle and right: enhancer activity changes (log2 FC) after mutating AP-1 or ETS instances at close (< 25 bp) or longer (> 50 bp) distance. Number of instances are shown. **** p-value < 0.0001 and ** < 0.01 (Wilcoxon rank-sum test). Box plots as in (B). **G)** Motif instances need to be analyzed within their cis-regulatory context. Motif syntax rules, such as motif combination, flanks and distance dictate the contribution of TF motif instances in enhancer sequences. Important motif instances will have a higher impact on enhancer activity when mutated.

Having trained a convolutional neural network to learn the motif syntax rules for *Drosophila* enhancers, we wanted to determine if the same type of rules also apply to human enhancers. Therefore, we generated simple linear models based on these rules to predict the contribution of individual motif instances in human enhancers. Specifically, these models consider the number of instances, the motif core and flanking sequence, and distance to other TF motifs (Fig 6E, S11B,C). Despite their simplicity, these models were able to predict motif-instance importance, with PCCs to experimentally assessed log2 fold-changes of 0.66 (P53), 0.61 (ETS), 0.59 (MAF) and 0.5 (AP-1), outperforming models based solely on PWM scores (Fig S11D). For most TFs, motif instances closer to an AP-1 or ETS motif were more important, suggesting that high cooperativity with these TFs is important in HCT116 enhancer sequences (Fig 6E, S11B). This was also observed between AP-1 and ETS motifs themselves, where mutation of either AP-1 or ETS instances had stronger impact in enhancer function if located at close (< 25 bp) rather than longer distances (> 50 bp) from each other (Fig 6F), and similarly between two AP-1 instances (Fig S11E). Altogether, these results confirm that the motif syntax rules derived for motif flanking sequence and inter-motif distances dictate the contribution of individual TF motif instances in human enhancers (Fig 6G). Determining how distinct instances of the same motif differentially contribute to enhancer activity could improve the ability to predict the functional impact of disease-associated variants, which typically affect only one motif instance.

### DeepSTARR designs synthetic enhancers with desired activities

Understanding how DNA sequence encodes enhancer activity should enable the design of synthetic enhancers with desired activity levels. We used DeepSTARR to computationally design new S2 cell developmental enhancers, by predicting enhancer activity for one billion random 249 bp DNA sequences that are not present in the *Drosophila* genome (see Methods). We then selected 249 of these sequences spanning different predicted activity levels and experimentally measured their enhancer activity by UMI-STARR-seq in S2 cells. The predicted activity of the synthetic sequences was highly accurate (PCC=0.62; Fig 7A) and DeepSTARR was able to design synthetic enhancers as strong as the strongest native S2 developmental enhancers (activity (fold- change over negative regions) ≈ 500; Supplementary Table 17).

**Figure 7.**
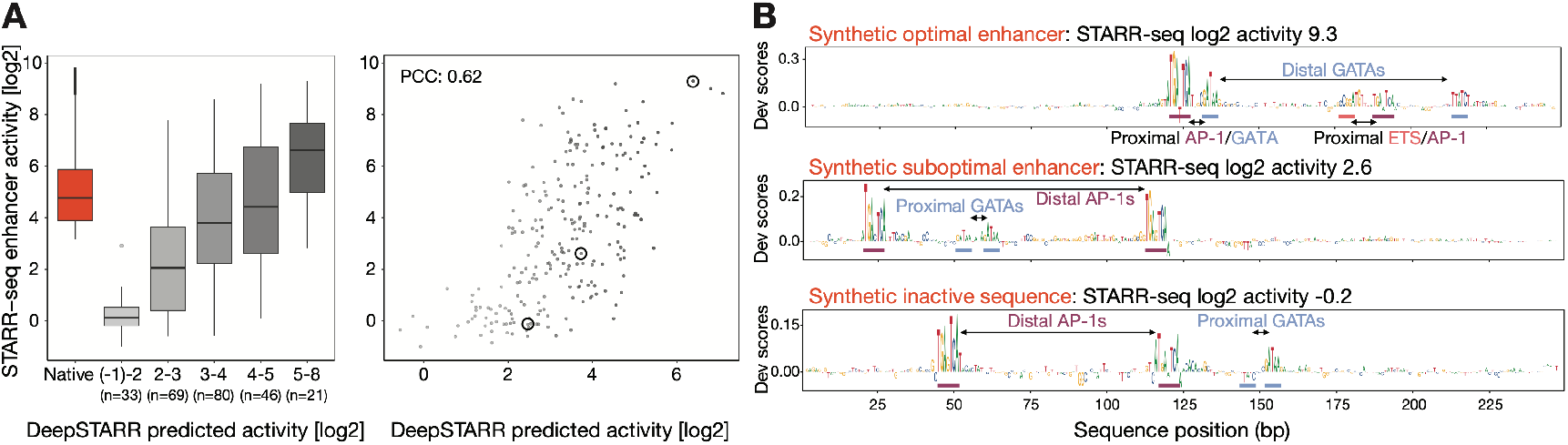
DeepSTARR designs synthetic enhancers using optimal sequence rules. **A)** Comparison between DeepSTARR predicted and experimentally measured enhancer activity (log2) for 249 synthetic sequences binned (left) or not (right). The ‘‘Native’’ category contains all *Drosophila* developmental enhancer sequences. The box plots mark the median, upper and lower quartiles and 1.5× interquartile range (whiskers); outliers are shown individually. The three synthetic sequences shown in (B) are highlighted. **B)** DeepSTARR nucleotide contribution scores for three synthetic sequences from (A) spanning different activity levels. Instances of GATA, AP-1 and ETS motifs are shown together with their observed distances (proximal or distal).

Inspection of the designed sequences suggested that their different activity levels correlated not only with motif composition but also the motif syntax (Fig 7B). For example, three different synthetic sequences, all containing two GATA and two AP-1 motifs, were predicted by DeepSTARR and validated experimentally to have very different activities (from 0.87 to 630). Interestingly, the strongest synthetic enhancer followed the optimal spacing rules predicted by DeepSTARR, such as distal GATA instances and proximal AP-1/GATA and ETS/AP-1 instances, whereas the other two synthetic sequences contained motifs in suboptimal syntax, such as distal AP-1 instances and proximal GATA instances (Fig 7B). This proof-of-concept experiment shows that the rules learned by DeepSTARR enable the *a priori* design of synthetic enhancers with desired activity levels.

## Discussion

Deciphering the rules governing the relationship between enhancer sequence and function – typically called the *cis-regulatory code* of enhancers – has remained a long-standing open problem. It has proved so challenging because methods to functionally characterize large numbers of enhancers have only become available a few years ago and also because the cis-regulatory code, unlike the protein-coding *genetic code*, follows complex and cell type-specific sequence-rules.

To dissect the relationship between enhancer sequence and activity for a single model cell type, we built a deep learning model, DeepSTARR, that accurately predicts enhancer activity for two different transcriptional programs directly from DNA sequence. DeepSTARR learned important TF motif types and higher-order syntax rules: different instances of the same TF motif are not functionally equivalent, and the differences are determined by motif flanks and inter-motif distances. These types of rules are also important in human enhancers and will be relevant to predict the impact of genetic variants linked to disease in the human genome.

The discovery that relatively rare sequence features can be important and predictive of enhancer activity is important and unexpected and highlights the potential of unbiased deep learning models that are not based on over-representation^47, 77^. The fact that motifs are often not arranged in optimal syntax agrees with previous work that suggested that suboptimal enhancers might have evolved to allow cell type specificity^12, 13^. Consistent with this interpretation, we observed optimized sequences of housekeeping enhancers that operate in all cell types.

Our results reveal an underappreciated property of enhancers: identical instances of the same TF motif with non-equivalent contributions to enhancer activity. Although the observation that only a small fraction of potential motifs throughout the genome is actually bound^21, 78, 79^ suggests that motif instances cannot all be equivalent, the non-equivalence of motif instances within the same enhancer is surprising. In fact, previous studies and computational models have typically considered different motif instances solely according to their PWM scores or even as equivalent^17, 26, 80^. The contribution of motif instances depended on high-order motif syntax rules such as inter-motif distances that are not captured by traditional PWM models and need to be modelled within the full enhancer sequence. This is in line with the recently reported limitations of PWM models for predicting the effects of noncoding variants on TF binding *in vitro*^81^ and improved performance of deep learning models for the prediction of motif instances bound *in vivo*^47, 57^. Together these results suggest that motif instances need to be analysed within their cis-regulatory context, which should improve our ability to predict and interpret the impact of disease-related sequence variants that typically affect individual motif instances.

The rules learned by DeepSTARR allowed the *de novo* design of synthetic S2 cell enhancers with desired activity levels, which not only demonstrates the validity of the model and its rules but also illustrates the power of this approach. Although libraries of synthetic elements have been used to explore enhancer structure^68^, it has remained impossible to build fully synthetic sequences with specific characteristics. It is interesting how these synthetic enhancers are of similar complexity as endogenous enhancers, e.g. in terms of motif number and diversity, and that a vast number of different sequences can have similar enhancer strengths, highlighting regulatory sequence flexibility and evolutionary opportunities. We expect that combining DeepSTARR with emerging algorithms that allow the direct generation of DNA sequences from deep learning models^54^ will provide unanticipated opportunities for the engineering of synthetic enhancers.

A next key challenge for the field will be to generalize such models from individual deeply characterized model cell lines to all cell types of an organism or even across species. This task is challenging because enhancers form the basis of differential gene transcription, and their activities are inherently cell-type specific. The underlying sequences and rules must therefore – by definition – also differ between cell types, at least to some extent. It is well known for example that enhancers that are active in different cell types or tissues contain different TF motifs^26, 80, 82^, which enables the binding of cell type-specific TFs. Therefore, it remains unclear how and to what extent cis-regulatory rules generalize or even apply universally.

We show here that differences between motif instances as well as the importance of motif flanks and distances generalize from *Drosophila* to human enhancers.

Unexpectedly, for AP-1 motifs, which we could assess in both species, the *Drosophila*- trained DeepSTARR model was able to predict the importance of AP-1 instances in human enhancers (PCC=0.42; Fig S12) and in both species ETS-AP-1 pairs synergize only at short distances but not at longer ones (mode 1; Fig S8C and Fig 6E,F, S11B). Ultimately, this demonstrates that although the specific rules vary between TF motif types and motif combinations, the types of rules as well as some specific rules apply more generally. Dissecting important types of rules in model cell lines together with the wealth of genomic data across many cell types (such as those from ENCODE) should unveil the gene-regulatory information in our genomes and a general cis-regulatory code.

## Methods

### UMI-STARR-seq

#### Cell culture

##### Drosophila S2 cells

Schneider 2 cells were grown in Schneider’s Drosophila Medium (Gibco; 21720-024) supplemented with 10% heat inactivated FBS (Sigma; F7524) at 27°C. Cells were passaged every 2-3 days.

##### Human HCT116 cells

Human HCT116 cells were cultured in DMEM (Gibco; 52100-047) supplemented with 10% heat inactivated FBS (Sigma; F7524) and 2mM L-Glutamine (Sigma; G7513) at 37°C in a 5% C02-enriched atmosphere. Cells were passaged every 2-3 days.

##### Electroporation

The MaxCyte-STX system was used for all electroporations. S2 cells were electroporated at a density of 50 x 10^7^ cells per 100µL and Sµg of DNA using the “Optimization 1” protocol. HCT116 cells were electroporated at a density of 1 x 10^7^ cells per 100µL and 20µg of DNA using the preset “HCT116” program.

#### UMI-STARR-seq experiments

##### Library cloning

*Drosophila* genome-wide libraries were generated by shearing genomic DNA from the sequenced *D.mel* strain (y; cn bw sp) to an average of 200 bp fragments. Inserts were cloned into the standard *Drosophila* STARR-seq vector^61^ containing either the DSCP or Rps12 core-promoters, and libraries grown in 6l of LB-Amp.

*Drosophila* and human oligo libraries were synthesized by Twist Bioscience including 249 bp enhancer sequence and adaptors for library cloning. Fragments from the *Drosophila* library were amplified (primers see Supplementary Table 1) and cloned into *Drosophila* STARR-seq vectors containing either the DSCP or Rps12 core-promoters using Gibson cloning (New England BioLabs; E2611S). The oligo library for human STARR-seq screens was amplified (primers see Supplementary Table 1) and cloned into the human STARR-seq plasmid with the ORI in place of the core promoter^83^. Libraries were grown in 2l LB-Amp.

All libraries were purified with Qiagen Plasmid *Plus* Giga Kit (cat. no. 12991).

##### Drosophila S2 cells

UMI-STARR-seq was performed as described previously^61, 62^. In brief, the screening libraries were generated from genomic DNA isolated of the sequenced *D.mel* strain (y; cn bw sp) or synthesized as oligo pools by Twist Bioscience (see above). We transfected 400 × 10^6 S2 cells total per replicate with 20 μg of the input library using the MaxCyte electroporation system. After 24 hr incubation, poly-A RNA was isolated and processed as described before^62^. Briefly: after reverse transcription and second strand synthesis a unique molecular identifier (UMI) was added to each transcript. This is followed by two nested PCR steps, each with primers that are specific to the reporter transcripts such that STARR-seq does not detect endogenous cellular RNAs.

##### Human HCT116 cells

STARR-seq was performed as described previously^61, 62, 83^. Screening libraries were generated from synthesized oligo pools by Twist Bioscience (see above). We transfected 80 × 10^6 HCT116 cells total per replicate with 160 µg of the input library using the MaxCyte electroporation system. After 6 hr incubation, poly-A RNA was isolated and further processed as described before^62^.

##### Illumina sequencing

Next-generation sequencing was performed at the VBCF NGS facility on an Illumina HiSeq 2500, NextSeq 550 or NovaSeq SP platform, following manufacturer’s protocol. Genome-wide UMI-STARR-seq screens were sequenced as paired-end 36 cycle runs (except the developmental input library, as paired-end 50 cycle runs) and Twist-oligo library screens were sequenced as paired-end 150 cycle runs, using standard Illumina i5 idexes as well as unique molecular identifiers (UMIs) at the i7 index.

#### Genome-wide UMI-STARR-seq data analysis

Paired-end genome-wide UMI-STARR-seq RNA and DNA input reads (36 bp; except the developmental input library that was 50 bp) were mapped to the *Drosophila* genome (dm3), excluding chromosomes U, Uextra, and the mitochondrial genome, using Bowtie v.1.2.2^84^. Mapping reads with up to three mismatches and a maximal insert size of 2 kb were kept. For paired-end RNA reads that mapped to the same positions, we collapsed those that have identical UMIs (10 bp, allowing one mismatch) to ensure the counting of unique reporter transcripts (Supplementary Table 2). We further computationally selected both RNA and input fragments of length 150-250 bp to only capture active sequences derived from short fragments. After processing the two biological replicates separately, we pooled both replicates for developmental and housekeeping screens for further analyses.

Peak calling was performed as described previously^61^. Peaks that had a hypergeometric p-value <= 0.001 and a corrected enrichment over input (corrected to the conservative lower bound of a 95% confidence interval) greater than 3 were defined as enhancers and resized to 249 bp (same length as used in oligo libraries) (Supplementary Table 3). Non-corrected enrichment over input was used as enhancer activity metric. Enhancers were classified as developmental or housekeeping based on the screen with the highest activity.

#### Oligo library UMI-STARR-seq data analysis

Oligo library UMI-STARR-seq RNA and DNA input reads (paired-end 150 bp) were mapped to a reference containing 249 bp long sequences containing both wildtype and mutated fragments from the *Drosophila* or human libraries using Bowtie v.1.2.2^84^. For the *Drosophila* library we demultiplexed reads by the i5 and i7 indexes and oligo identity. Mapping reads with the correct length, strand and with no mismatches (to identify all sequence variants) were kept. Both DNA and RNA reads were collapsed by UMIs (10 bp) as above (Supplementary Table 2).

We excluded oligos with less than 10 reads in any of the input replicates and added one read pseudocount to oligos with zero RNA counts. The enhancer activity of each oligo in each screen was calculated as the log2 fold-change over input, using all replicates, with DESeq2^85^. We used the counts of wildtype negative regions in each library as scaling factors between samples. This normalization only changes the position of the zero and consequently does not affect the calculation of log2 fold-changes between different sequences or the p-values for the statistical tests used.

### Deep Learning

#### Data preparation

We selected all windows at the summit of developmental and housekeeping enhancers, in addition to three windows on either side of the regions (stride 100 bp). The remaining part of the genome was binned into 249 bp windows with a stride of 100 bp, excluding chromosomes U, Uextra, and the mitochondrial genome. We only included bins with more than five reads in the input and at least one read in the RNA of both developmental and housekeeping screens. To have a diversity of inactive sequences, we selected (1) 20,000 random bins overlapping accessible regions in different *Drosophila* cell types (S2, kc167 and OSC^61, 86^) and embryogenesis stages^87^, as well as all bins overlapping (2) enhancers from different *Drosophila* cell types (OSC and BG3^26^) and (3) inducible enhancers in S2 cells for two different stimuli (ecdysone^88^ and Wnt signaling^89^). Lastly, we added 59,081 random windows with a range of enhancer activity levels. We augmented our dataset by adding the reverse complement of each original sequence, with the same output, ending up with 242,026 examples (484,052 post-augmentation). Sequences from the first (40,570; 8.4%) and second half of chr2R (41,186; 8.5%) were held out for validation and testing, respectively.

#### DeepSTARR model architecture and training

DeepSTARR was designed as a multi-task convolutional neural network (CNN) that uses one-hot encoded 249 bp long DNA sequence (A=[1,0,0,0], C=[0,1,0,0], G=[0,0,1,0], T=[0,0,0,1]) to predict both its developmental and housekeeping enhancer activities (Fig 1C). We adapted the Basset CNN architecture^45^ and built DeepSTARR with four 1D convolutional layers (filters=246,60,60,120; size=7,3,5,3), each followed by batch normalization, a ReLU non-linearity, and max-pooling (size=2). After the convolutional layers there are two fully connected layers, each with 256 neurons and followed by batch normalization, a ReLU non-linearity, and dropout where the fraction is 0.4. The final layer mapped to both developmental and housekeeping outputs. Hyperparameters were manually adjusted to yield best performance on the validation set. The model was implemented and trained in Keras (v.2.2.4^90^) (with TensorFlow v.1.14.0^91^) using the Adam optimizer^92^ (learning rate = 0.002), mean squared error (MSE) as loss function, a batch size of 128, and early stopping with patience of ten epochs. Model training, hyperparameter tuning and performance evaluation were performed on different sets of genomic regions in distinct chromosomes.

#### Performance evaluation

The performance of the model was evaluated separately for developmental and housekeeping predictions on the held-out test sequences. We used the Pearson correlation coefficient (PCC) across all bins for a quantitative genome-wide evaluation and the area under the precision-recall curve (AUPRC; calculated using pr.*curve* from R package *PRROC* v.1.3.1^92^) for enhancer classification (enhancers vs. 2,685 negative control regions from the test set).

To test the robustness of the model, we trained 1,000 DeepSTARR models with the same set of hyperparameters and compared their performance. This accounted for the stochastic heterogeneity due to the random initialized weights in the neural network.

#### Prediction on full *Drosophila* genome

We extracted 249 bp sequences tiled across the *Drosophila* dm3 genome with a stride of 20 bp using ‘‘bedtools makewindows’’ (parameters -w 249 -s 20’) and ‘‘bedtools getfasta”^93^. We next predicted the developmental and housekeeping enhancer activity of each genomic window with DeepSTARR and averaged these per nucleotide to obtain genome-wide coverage. The DeepSTARR predicted coverage tracks are shown as examples in Fig 1B and S1A,B and are available at https://genome.ucsc.edu/s/bernardo.almeida/DeepSTARR_manuscript.

#### Models for comparison

The performance of DeepSTARR in the test set sequences was compared with two different methods: (1) a gapped k-mer support vector machine (gkm-SVM)^39^ and (2) a lasso regression model based on TF motif counts (Fig S1D).

(1) We used a 10-fold cross-validation scheme to train a developmental and a housekeeping gkm-SVM model to classify 249 bp DNA sequences into enhancers. Training was performed using developmental or housekeeping enhancers and a set of 21,463 negative control regions from the training set. The gkm-SVMs were done using LS-GKM^94^ and the following parameters: (dev) *gkmtrain -t 0 -l 8 -k 5 -x 10*; (hk) *gkmtrain -t 0 -l 11 - k 7 -x 10*. We used the resulting support vectors of each trained model to score the DNA sequences of the test set by running *gkmpredict* and used these scores for the PCC and AUPRC analysis.
(2) We trained lasso regression models for developmental and housekeeping enhancer activity using the counts of 6,502 known TF motifs (see “Reference compendium of non-redundant TF motifs” below) as features across 40,000 random selected bins from the training set. Motif counts were calculated using the *matchMotifs* function from R package *motifmatchr* (v.1.4.0^95^) with the following parameters: *genome = “BSgenome.Dmelanogaster.UCSC.dm3”, p.cutoff = 5e^-04^, bg=“even”*. The model was trained using the optimal *lambda* retrieved from 10-fold cross-validation and the *glmnet* function from R package *glmnet* (v.2.0-16^96^).

#### Nucleotide contribution scores

We used DeepExplainer (the DeepSHAP implementation of DeepLIFT, see refs. ^55, 63, 64^; update from https://github.com/AvantiShri/shap/blob/master/shap/explainers/deep/deep_tf.py) to compute contribution scores for all nucleotides in all sequences in respect to either developmental or housekeeping enhancer activity. We used 100 dinucleotide-shuffled versions of each input sequence as reference sequences. For each sequence, the obtained hypothetical importance scores were multiplied by the one-hot encoded matrix of the sequences to derive the final nucleotide contribution scores, which were visualized using the *ggseqlogo* function from R package *ggseqlogo* (v.0.1^97^).

#### Motif discovery using TF–Modisco

To consolidate motifs, we ran TF–Modisco (v.0.5.12.0^56^) on the nucleotide contribution scores for each enhancer type separately using all developmental or housekeeping enhancers (Fig 2B). We specified the following parameters: *sliding*_window_size=15, *flank*_size=5, max_seqlets_per_*metacluster*=50000 and *TfModiscoSeqletsToPatternsFactory(trim_to_window_size=15, initial_flank_to_add=5)*.

### Reference compendium of non-redundant TF motifs

#### Reference compendium of non-redundant TF motifs

6,502 TF motif models were obtained from iRegulon (http://iregulon.aertslab.org/collections.html ^98^) covering the following databases: Bergman (version 1.1^99^), CIS-BP (version 1.02^100^), FlyFactorSurvey (2010^101^), HOMER (2010^102^), JASPAR (version 5.0_ALPHA^103^), Stark (2007^104^) and iDMMPMM (2009^105^). We systematically collapsed redundant motifs by similarity by a previously described approach^76^. Specifically, we computed the distances between all motif pairs using TOMTOM^106^ and performed hierarchical clustering using Pearson correlation as the distance metric and complete linkage using the *hclust* R function. The tree was cut at height 0.8, resulting in 901 non-redundant motif clusters that were manually annotated (Fig S2A-E). Clustering of motifs from each cluster and their logos were visualized using the *motifStack* R package (v.1.26.0^107^). The code and TF motif compendium are available from https://github.com/bernardo-de-almeida/motif-clustering.

#### TF motif enrichment analyses in developmental and housekeeping enhancers

We tested the enrichment of each TF motif in developmental or housekeeping enhancers over negative genomic regions (Fig S2F,G, Supplementary Table 4). Counts for each motif in each sequence were calculated using the *matchMotifs* function from R package *motifmatchr* (v.1.4.0^95^) with the following parameters: *genome = “BSgenome.Dmelanogaster.UCSC.dm3”, p.cutoff = 1e^-04^, bg=“genome”*. For each enhancer type, we assessed the differential distribution of each motif between the enhancers and negative regions by two-sided Fisher’s exact test. Obtained P-values were corrected for multiple testing by Benjamini-Hochberg procedure and considered significant if FDR :: 0.05. To remove motif redundancy, only the most significant TF motif per motif cluster was shown.

### TF motif mutagenesis in *Drosophila* S2 enhancers

#### Oligo library design

##### Selection of enhancer regions

A comprehensive library of 5,082 wildtype enhancer sequences in *D. melanogaster* S2 cells was compiled by selecting previously published developmental^61^, housekeeping^40^ and inducible (ecdysone^88^ and Wnt signaling^89^) enhancers. 249 bp sequences centered on the enhancers’ summit in both forward and reverse orientation were retrieved. We added 524 249-bp negative genomic regions in both orientations as controls (Supplementary Table 5).

##### Generation of TF motif mutations

We selected four predicted developmental motifs (GATA, AP-1, twist, Trl), three predicted housekeeping motifs (Dref, Ohler1, Ohler6) and three control motifs (length-matched random motifs to control for enhancer-sequence perturbation). For each motif type, we mapped all instances using string-matching (GATA: GATAA; AP-1: TGA.TCA; twist: CATCTG/CATATG; Trl: GAGAG; Dref: ATCGAT; Ohler1: GTGTGACC; Ohler6: AAAATACCA; control: TAGG, GGGCT, CCTTA) in 2,194 enhancers (both motif orientations) and mutated all instances both simultaneously and each instance individually to a motif shuffled variant (Supplementary Table 5; Fig S3A). Each instance for a given motif was mutated always to the same shuffled variant to allow the comparison of effects between instances of the same motif type. We designed motif-mutant sequences for each enhancer only for the orientation with the strongest wildtype enhancer activity. In addition, for each motif type we repeated mutations with two other different shuffled variants in 50 enhancers to control for the impact of the selected shuffled variant (Supplementary Table 5; Fig S3C).

#### Enhancers with swapped GATA motif flanks

We selected 100 developmental enhancers from above that contain 2 GATA instances (*inst1* and *inst2*) with different importance as predicted by DeepSTARR and swapped the flanking nucleotides (both 2 bp and 5 bp separately) between both instances (Fig 4C, S7B). For each enhancer, we designed sequences where the flanks of *inst1* were replaced by the flanks of *inst2* and vice versa, resulting in sequences where both the two GATA instances contained either the flanks of *inst1* or the flanks of *inst2*. In addition, when replacing the flanks of *inst1* by the flanks of *inst2*, we also mutated *inst2* to assess how the flanks of *inst2* affected the contribution of *inst1*. The opposite was also done, with the flanks of *inst2* being replaced by the flanks of *inst1* together with mutation of *inst1*. The mutated sequences are listed in Supplementary Table 5. 47 active enhancers contained one strong and one weak GATA instances (≥ 2-fold difference as assessed afterwards by mutagenesis) were used for the analyses in Fig 4C and S7B (Supplementary Table 11).

##### Designing of synthetic S2 developmental enhancers

1 billion random 249 bp DNA sequences were generated in *bash* with the following code: *cat /dev/urandom | tr -dc ’ACGT’ | fold -w 249 | head -n 1000000000*. Bowtie v.1.2.2 ^84^ was used to remove sequences that exist in the *D. melanogaster* genome, which were none. The developmental enhancer activity of these sequences was predicted using DeepSTARR and 249 sequences spanning different activity levels were selected for the oligo library (Supplementary Table 5 and 17).

#### Oligo library synthesis and UMI-STARR-seq

The *Drosophila* enhancers’ motif mutagenesis oligo library contained wildtype (both orientations) and motif-mutant enhancers, enhancers with swapped GATA motif flanks and synthetic enhancer sequences (Supplementary Table 5). All sequences were designed using the dm3 genome version. The enhancer sequences spanned 249 bp total, flanked by the Illumina i5 (25 bp; 5 ′ -TCCCTACACGACGCTCTTCCGATCT) and i7 (26 bp; 5 ′ AGATCGGAAGAGCACACGTCTGAACT) adaptor sequences upstream and downstream, respectively, serving as constant linkers for amplification and cloning. The resulting 21,758-plex 300-mer oligonucleotide library was synthesized by Twist Biosciences Inc. UMI-STARR-seq using this oligo library was performed (“UMI-STARR-seq experiments”) and analyzed (“Oligo library UMI-STARR-seq data analysis”) as described above. We performed three independent replicates for developmental and housekeeping screens (correlation PCC=0.93-0.98; Fig S3B).

#### TF motif mutation analysis and equivalency

From the candidate 249 bp enhancers, we identified 855 active developmental and 905 active housekeeping *Drosophila* enhancers (log2 wildtype activity in oligo UMI-STARR- seq >= 3.15 and 2.51, respectively; the strongest negative region in each screen) that we used in the subsequent TF motif mutation analyses. The impact of mutating all instances of a TF motif type simultaneously or each instance individually was measured as the log2 fold-change enhancer activity between the respective mutant and wildtype sequences (Supplementary Table 6 and 8). This was done separately for developmental and housekeeping enhancer activities.

Motif non-equivalency across all enhancers (Fig 3B, S5B,D) or within the same enhancer (Fig 3A,C) was assessed by comparing the impact of mutating individual instances of the same TF motif, i.e. the log2 fold-changes of each instance (Supplementary Table 8). For the comparison between instances in the same enhancer, only enhancers that require the TF motif (> 2-fold reduction in activity after mutating all instances) and contain two or more instances were used. Motif instances with >2-fold different contributions in the same enhancer were considered as non-equivalent. The same comparison across enhancers or within the same enhancer was performed for the three control motifs.

### Motif syntax features

#### DeepSTARR predicted global importance of motif types and comparison with motif enrichment

To quantify the global importance of all known TF motifs to enhancer activity *in silico* (see ref. ^58^), we embedded each motif from the 6,502 TF motif compendium at five different locations and in both strands in 100 random backbone DNA sequences and predicted their developmental and housekeeping enhancer activity with DeepSTARR. The 249 bp backbone sequences were generated by sampling the base at each position with equal probability. The five different locations were the same for all motifs, centered at positions 25, 75, 125 (middle of the 249 bp oligo), 175 and 225. For each motif, we used the sequence corresponding to the highest affinity according to the annotated PWM models. The average activity across the different locations per backbone was divided by the backbone initial activity to get the predicted increase in enhancer activity per TF motif.

The resultant log2 fold-change was averaged across all 100 backbones to derive the final global importance of each TF motif.

The global motif importance predicted by DeepSTARR was compared with the enrichment of TF motifs at developmental and housekeeping enhancers, measured as the two-sided Fisher’s exact test log2 odds ratio (described in “TF motif enrichment analyses in developmental and housekeeping enhancers”) (Fig 2D, Supplementary Table 7). To remove motif redundancy, only the TF motif with the strongest predicted global importance or the strongest motif enrichment per motif cluster are shown in Fig 2D.

#### DeepSTARR predictions for the contribution of motif instances

We used two complementary approaches to measure the predicted contribution of each motif instance by DeepSTARR.

First, we measured the predicted importance of all string-matched instances of each TF motif type in 9,074 developmental enhancers, 6,369 housekeeping enhancers or 26,938 negative genomic regions (Fig S5A,C; Supplementary Table 9). The predicted importance of an instance was calculated as the average developmental or housekeeping DeepSTARR contribution scores over all its nucleotides. These scores represent the global contribution of motif instances captured by the model and were used for the analyses of figures: 4A,B, S5A,C, S7A.

Second, to compare with the experimentally derived motif importance through motif mutagenesis, we used DeepSTARR to predict the log2 fold-change between wildtype and the motif-mutant enhancer sequences included in the oligo library for all instances of the different motif types (Fig 3B,D, S6). This was done by calculating the log2 fold-change between the predicted activity of the wildtype and respective motif-mutant sequences. Since the experimentally derived importance can be dependent on the shuffled mutant variant selected, this provides a more direct evaluation of the capability of DeepSTARR to predict the importance of a motif instance assessed by experimental mutagenesis.

#### Scoring of TF motif instances with PWM motif scores

To assess how the PWM motif models predict the importance of a motif instance, we scored the wildtype sequence of each mutated motif instance (extended 10 nucleotides on each flank to account for the flanking sequence) with the PWM models of the selected TF motifs (Supplementary Table 10). We used the *matchMotifs* function from R package *motifmatchr* (v.1.4.0; *genome = “BSgenome.Dmelanogaster.UCSC.dm3”, bg=“even”*^95^) with a p-value cutoff of 1 to retrieve the PWM scores of all sequences. These PWM scores were compared with the experimental log2 fold-changes using Pearson correlation (Fig 3D).

We tested different PWM models for each TF motif if available and reported always the one with the best correlation (Supplementary Table 10).

#### Correlation between motif importance and motif flanks

String-matched motif instances of each TF were sorted by their predicted (DeepSTARR) or experimentally derived (minus signed (-) mutation log2 fold-change) importance. Their 5 flanking nucleotides were shown using heatmaps and the importance of each nucleotide at each flanking position summarized using box plots (Fig 4A, S7A). Significant differences between the four nucleotides per position were assessed through Welch One-Way ANOVA test followed by FDR multiple testing correction. The motif logos represent the frequency of each nucleotide at each position among the top 90^th^ percentile instances and were compared with the logos of existing PWM models (Fig 4B).

#### *In silico* motif distance preferences

Two consensus TF motifs were embedded in 60 random backbone 249 bp DNA sequences, *MotifA* in the center and *MotifB* at a range of distances (*d*) from *MotifA*, both up- and downstream (Fig 5A, S8). Backbone sequences were generated by sampling the base at each position with equal probability. DeepSTARR was used to predict the developmental or housekeeping activity of the backbone synthetic sequences (1) without any motif (*b*), (2) only with *MotifA* in the center (*A*), (3) only with *MotifB d*-bases up- or downstream (*B*) and (4) with both *MotifA* and *MotifB* (*AB*). The cooperativity between *MotifA* and *MotifB* at each distance *d* was then defined as the fold-change between *AB* and (*b* + (*A*-*b*) + (*B*-*b*) = *A*+*B*-*b*), where a value of 1 means an additive effect or no synergy between the motifs, and a value higher than 1 means positive synergy. The median of fold-changes across the 60 backbones was used as the final cooperativity scores. This analysis was performed for all motif pair combinations of AP-1, SREBP, GATA, Trl, twist and ETS motifs for developmental enhancer activity, and Dref, Ohler1 and Ohler6 for housekeeping enhancer activity in both strand orientations. Pairs with a negative control motif (GGGCT) were also included.

#### Enrichment of motif pairs at different distances in genomic enhancers

We obtained the positions of the different TF motif instances across all 9,074 developmental enhancers, 6,369 housekeeping enhancers and 26,938 negative genomic regions as described above (“Correlation between motif number and enhancer activity”). To compute whether *MotifA* is located within a certain distance (bins: 0-25, 25-50, 50-75, 75-100, 100-125, 125-150, 150-250 bp) of *MotifB* more/less frequently in enhancers than in negative sequences, we counted the number of times a *MotifA* instance is at each distance bin to a *MotifB* instance in enhancers and in negative sequences. The enrichment or depletion of motif pairs at each bin was tested with two-sided Fisher’s exact test and the log2 odds ratio used as metric. Obtained P-values were corrected for multiple testing by Benjamini-Hochberg procedure and considered significant if FDR :: 0.05. We performed this analysis separately for all developmental motif pairs in developmental enhancers and all housekeeping motif pairs in housekeeping enhancers (Fig 5B, S9A,C,D).

#### Association between motif pair distances and enhancer activity

We obtained the positions of the different TF motif instances across all 9,074 developmental enhancers, 6,369 housekeeping enhancers and 26,938 negative genomic regions as described above (“Correlation between motif number and enhancer activity”). For each pair of motif instances at each distance bin (0-25, 25-50, 50-75, 75-100, 100- 125, 125-150, 150-250 bp), we tested the association between enhancer activity and the presence of the pair at the respective distance bin using a multiple linear regression, including as independent variables the number of instances for the different developmental or housekeeping TF motif types. The linear model coefficient was used as metric and considered significant if the FDR-corrected p-values :: 0.05. We performed this analysis separately for all developmental motif pairs in developmental enhancers and all housekeeping motif pairs in housekeeping enhancers (Fig 5B, S9B-D).

#### Validation of motif distance preferences by motif mutagenesis

To test how the importance of GATA and AP-1 instances associate with the absolute distance *d* to a second GATA instance, we compared the log2 fold-change in enhancer activity after mutating individual GATA (Fig 5D) or AP-1 (Fig 5E) instances at close (< 25 bp; n=14 and 29, respectively) or longer (> 50 bp; n=129 and 38) distance to a second GATA instance. Only pairs of non-overlapping motif instances were used. A Wilcoxon rank-sum test was used to test this association.

### TF motif mutagenesis in human HCT116 enhancers

#### TF motif enrichment

We characterized the motif composition of 5,891 strong STARR-seq enhancers in human HCT116 cells^83^ using the 501 bp sequence centered on the summit. We generated 5,891 negative GC-matched genomic regions using the *genNullSeqs* function from R package *gkmSVM*^108^. 1,689 TF motif PWM models and respective motif clustering information were retrieved from Vierstra et al.,^76^ covering the following databases: JASPAR (2018), Taipale HT-SELEX (2013) and HOCOMOCO (version 11). Counts for each motif in each 501 bp enhancer and negative sequence were calculated using the *matchMotifs* function from R package *motifmatchr* (v.1.4.0^95^) with the following parameters: *genome = “BSgenome.Hsapiens.UCSC.hg19”, p.cutoff = 1e^-04^, bg=“genome”*. We assessed the differential distribution of each motif between the enhancers and negative regions by two-sided Fisher’s exact test. We selected the nine TF motifs with the strongest enrichment in enhancers: AP-1, P53, MAF, CREB1, ETS, EGR1, MECP2, E2F1 and Ebox/MYC (Supplementary Table 12).

#### TF motif mutagenesis oligo library design and synthesis

##### Generation of TF motif mutations

For UMI-STARR-seq of wild type and mutant enhancers, we selected 3,200 enhancer candidates, defining short 249 bp windows (the limits of oligo synthesis), and mapped the position of all instances of the nine TF motif types in these candidates using the *matchMotifs* function from R package *motifmatchr* (v.1.4.0^95^) with the following parameters: *genome = “BSgenome.Hsapiens.UCSC.hg19”, p.cutoff = 5e^-04^, bg=“genome”*. Overlapping instances (minimum 70%) for the same TF motif were collapsed. We also mapped all instances of four control motifs (length-matched random motifs to control for enhancer-sequence perturbation) using string-matching. We then designed enhancer variants with all instances of each motif type mutated simultaneously or individually to a motif shuffled variant (Supplementary Table 13; Fig S10A). Each instance for a given motif was mutated always to the same shuffled variant to allow the comparison of effects between motif instances. We designed motif-mutant sequences for each enhancer only for the orientation with the strongest activity in the genome-wide STARR-seq. In addition, for each motif type we repeated mutations with two other different shuffled variants in 50 enhancers to control for the impact of the selected shuffled variant (Supplementary Table 13; Fig S10F).

##### Oligo library synthesis and UMI-STARR-seq

The final human enhancers’ motif mutagenesis library contained 3,200 wildtype and 18,780 motif-mutant enhancer sequences that we combined with 920 249-bp negative genomic regions as controls (Supplementary Table 13). All sequences were designed using the hg19 genome version. Apart from the specific sequences, this human motif mutagenesis library exhibits the same specifications as the *Drosophila* library and was also synthesized by Twist Biosciences Inc. UMI-STARR-seq using this oligo library was performed (“UMI-STARR-seq experiments”) and analyzed (“Oligo library UMI-STARR-seq data analysis”) as described above. We performed two independent replicates (correlation PCC=0.99; Fig S10B).

#### TF motif mutation analysis

From the 3,200 designed candidate 249 bp enhancers, we identified 1,083 active short human enhancers (log2 wildtype activity in oligo UMI-STARR-seq >= 2.03, the strongest negative region; Fig S10C) that we used in the subsequent TF motif analyses. The impact of mutating all instances of a TF motif type simultaneously or each instance individually was calculated as the log2 fold-change enhancer activity between the respective mutant and wildtype sequences (Fig S10D,E, S11A; Supplementary Table 14 and 15). Motif non-equivalency across all enhancers (Fig S11A) or within the same enhancer (Fig 6B,C) was assessed as in the *Drosophila* enhancers.

#### Validation of important TF motif instances with genomic DNase I footprinting data

We compared the importance of individual motif instances with genomic DNase I footprinting data of RKO cells (another human colon cancer cell line; https://www.vierstra.org/resources/dgf ^76^), as a surrogate for TF occupancy (Fig 6D). Footprints detected at different FPR adjusted p-value thresholds and coverage tracks with observed and expected cleavage counts were downloaded from https://resources.altius.org/~jvierstra/projects/footprinting.2020/per.dataset/h.RKO-DS40362/, in hg38 coordinates. All coordinates were converted to hg19 coordinates using the *UCSC liftOver* tool ^109^ and the *hg38ToHg19.over.chain* chain file (https://hgdownload.soe.ucsc.edu/goldenPath/hg38/liftOver/hg38ToHg19.over.chain.gz). For each TF motif type, a Wilcoxon rank-sum test was used to determine whether the mutation log2 fold-change of instances overlapping TF footprints (FPR threshold of 0.001) is significantly greater or less than the one of instances not overlapping footprints. Only instances within HCT116-accessible enhancers were used in the analysis. Enhancers were defined as accessible if they overlap any of the DNase-seq peaks from the following ENCODE^75^ identifiers (hg19 coordinates) (https://www.encodeproject.org/): ENCFF001SQU, ENCFF001WIJ, ENCFF001WIK, ENCFF175RBN, ENCFF228YKV, ENCFF851NWR, ENCFF927AHJ, ENCFF945KJN and ENCFF360XGA.

#### Association between motif syntax rules and the contribution of TF motif instances

For each TF motif type, we built a multiple linear regression model to predict the contribution of its individual instances (log2 fold-changes) using as covariates the number of instances of the respective motif type in the enhancer, the motif core (defined as the nucleotides included in each TF motif PWM model) and flanking nucleotides (5 bp on each side), and the distance to all other TF motifs (close: < 25 bp; intermediate: ≥ 25 bp and :: 50 bp; distal: >50 bp) (Fig 6E, S11B-D). Only motif instances that start after position 5 and end before position 245 of the 249 bp oligos were used, in order to be able to retrieve their 5 bp flanking sequences. In addition, for the motif distance analyses only non-overlapping motif pairs were used. All models were built using the *Caret* R package (v. 6.0-80^110^) and 10-fold cross-validation. Predictions for each held-out test sets were used to compare with the observed log2 fold-changes and assess model performance. The linear model coefficients and respective FDR-corrected p-values were used as metrics of importance for each feature (Fig 6E, S11B). For each TF motif type, we compared the main regression model with a simple linear model only using the PWM scores as covariate (Fig S11D).

#### DeepSTARR prediction of the importance of AP-1 instances in human enhancers

We used the DeepSTARR model trained in *Drosophila* S2 enhancers to predict the importance of AP-1 instances in human HCT116 enhancers. This was done by predicting the activity of the wildtype and motif-mutant enhancer sequences included in the human oligo library for all AP-1 instances and further calculating the log2 fold-change. This predicted log2 fold-change was compared with the experimentally measured log2 fold-change and its association assessed through Pearson correlation (Fig S12; Supplementary Table 16).

### Statistics and data visualization

All statistical calculations and graphical displays have been performed in R statistical computing environment (v.3.5.1^111^) and using the R package *ggplot2* (v.3.2.1^112^). Coverage data tracks have been visualized in the UCSC Genome Browser^113^ and used to create displays of representative genomic loci. In all box plots, the central line denotes the median, the box encompasses 25th to 75th percentile (interquartile range) and the whiskers extend to 1.5× interquartile range.

### Data availability

The raw sequencing data are available from GEO (https://www.ncbi.nlm.nih.gov/geo/) under accession number GSE183939. Data used to train and evaluate the DeepSTARR model as well as the final pre-trained model are found on zenodo at https://doi.org/10.5281/zenodo.5502060. We also plan to release the pre-trained DeepSTARR model in the Kipoi model repository^114^. Genome browser tracks showing genome-wide UMI-STARR-seq and DeepSTARR predictions in *Drosophila* S2 cells, together with the enhancers used for mutagenesis, mutated motif instances and respective log2 fold-changes in enhancer activity, are available at https://genome.ucsc.edu/s/bernardo.almeida/DeepSTARR_manuscript. TF motif models were obtained from iRegulon (http://iregulon.aertslab.org/collections.html ^98^). DNase-seq data in *Drosophila* S2 cells were obtained from ref.^61^. Genomic DNase I footprinting data of RKO cells were downloaded from https://resources.altius.org/~jvierstra/projects/footprinting.2020/per.dataset/h.RKO-DS40362/. HCT116 DNase-seq, H3K27ac and H3K4me1 data were obtained from ENCODE^75^ (https://www.encodeproject.org/) and ATAC-seq data from ref.^74^.

### Code availability

Code used to process the genome-wide and oligo UMI-STARR-seq data and train DeepSTARR, as well as to predict the enhancer activity for new DNA sequences is available on GitHub (https://github.com/bernardo-de-almeida/DeepSTARR). The code and TF motif compendium are available from https://github.com/bernardo-de-almeida/motif-clustering.

## Acknowledgements

The authors thank Angela Andersen (Life Science Editors), Vincent Loubiere and Franziska Lorbeer (IMP) for comments on the manuscript, and Gert Hulselmans and Stein Aerts (KU Leuven) for sharing the TF motif PWM collection. Deep sequencing was performed at the Vienna Biocenter Core Facilities GmbH. Research in the Stark group is supported by the European Research Council (ERC) under the European Union’s Horizon 2020 research and innovation programme (grant agreement no. 647320) and by the Austrian Science Fund (FWF, F4303-B09). Basic research at the IMP is supported by Boehringer Ingelheim GmbH and the Austrian Research Promotion Agency (FFG).

## Author Contributions

B.P.d.A., F.R. and A.S. conceived the project. F.R. and M.P. performed all experiments. B.P.d.A. performed all computational analyses. B.P.d.A., F.R. and A.S. interpreted the data and wrote the manuscript. A.S. supervised the project.

## Supplementary Figures

**Supplementary Figure 1.**
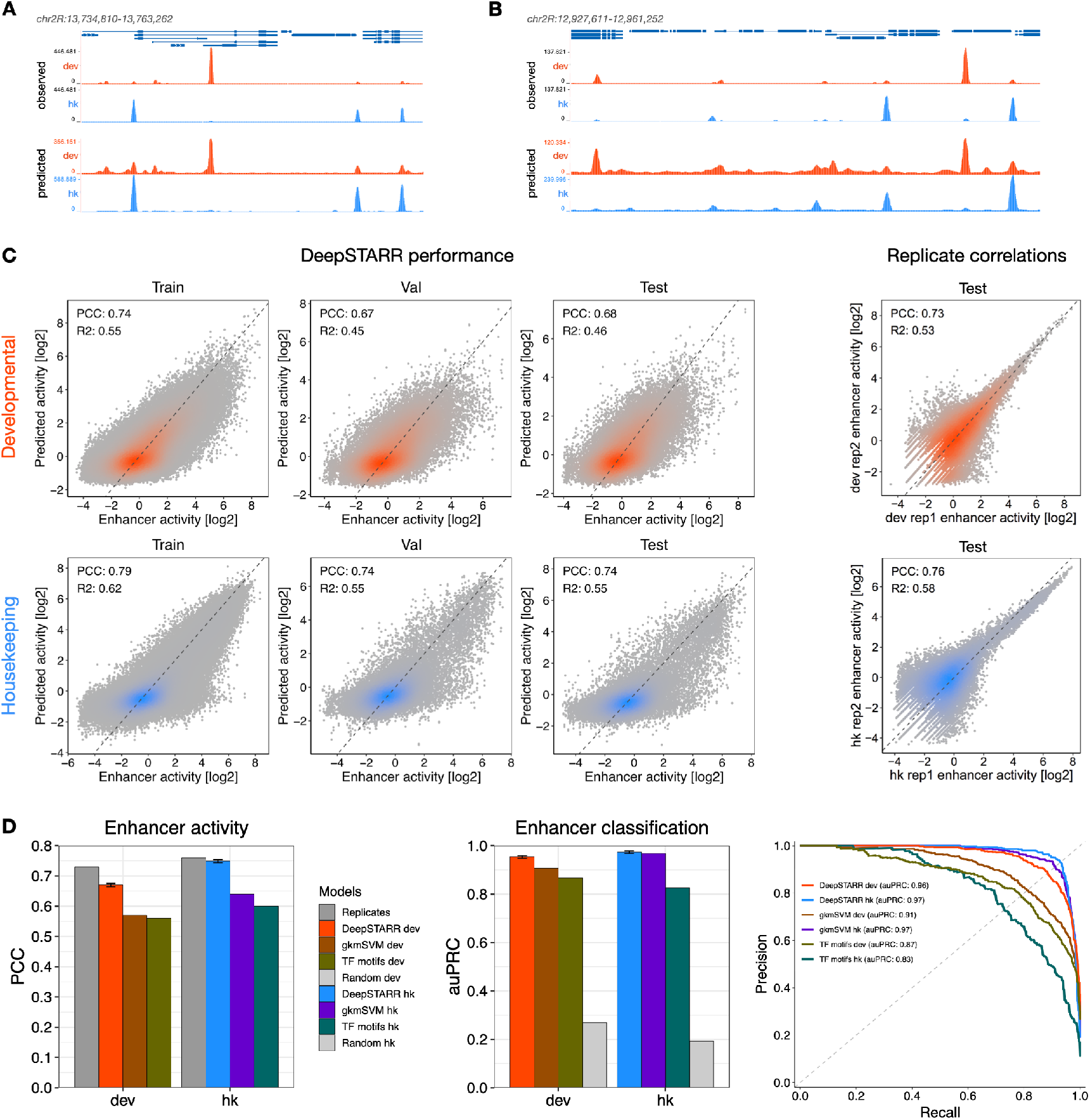
Additional performance evaluation of DeepSTARR predictions. **A-B)** DeepSTARR predicts enhancer activity genome-wide. Genome browser screenshot depicting UMI-STARR-seq observed (top) and predicted (bottom) profiles for both promoters (development, red; housekeeping, blue) for two loci located on held-out test chromosome 2R. **C)** DeepSTARR predicts enhancer activity quantitatively. Left: Scatter plots of predicted vs. observed developmental (top) and housekeeping (bottom) enhancer activity signal across all DNA sequences in the train, validation and test set chromosomes. Right: Scatter plots of developmental (top) and housekeeping (bottom) enhancer activity signal between two biological replicates across all DNA sequences in the test set chromosome. Color reflects point density. The PCC is denoted for each comparison. **D)** DeepSTARR performed better than methods based on known TF motifs or unbiased k-mers. Left: Comparison of different models for predicting enhancer activity. Bar-plots with the PCC between observed and predicted activities for both developmental and housekeeping enhancer types across all DNA sequences in the test set chromosome. PCC between replicates is also shown. Middle: Bar-plots with the auPRC for the classification of enhancer sequences from the test set for the different models, compared with the expected by a random model. Right: precision-recall curve for the different models on test data. Error bars represent the 5^th^ and 95^th^ percentile of the performance of 1000 DeepSTARR models. PCC: Pearson correlation coefficient, R2: R-squared, auPRC: area under precision-recall curve.

**Supplementary Figure 2.**
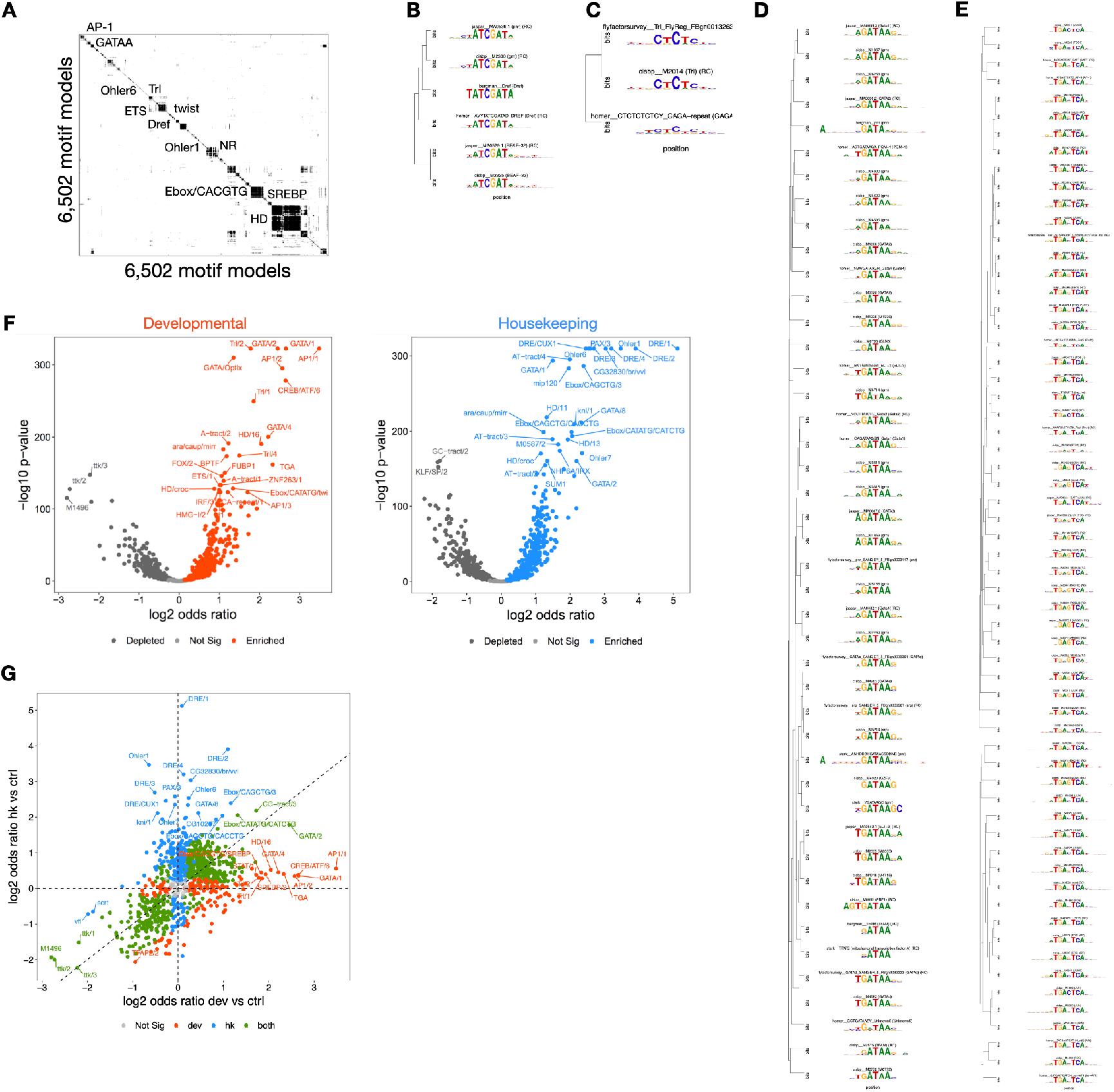
Developmental and housekeeping enhancers are enriched in different TF motifs. **A)** Hierarchically clustered heat map of the pairwise similarity scores between 6,502 TF motifs. The cluster dendrogram was cut at height 0.8, resulting in 901 non-redundant motif clusters that were manually annotated. **B-E)** Exemplar TF motif clusters. **F)** Enrichment of TF motifs in developmental (left) and housekeeping (right) enhancers over negative genomic regions. Log2 Fisher’s odds ratio compared with significance (-log10 p-value) for the most significant TF motif per motif cluster, to remove motif redundancy. Motifs significantly (FDR<0.05) enriched or depleted are highlighted. **G)** Scatter plot comparing the motif enrichment (log2 odds ratio) in developmental and housekeeping enhancers. To remove motif redundancy, only the most significant TF motif per motif cluster was shown. Motifs significantly (FDR<0.05) enriched or depleted in each or both enhancer types are highlighted.

**Supplementary Figure 3.**
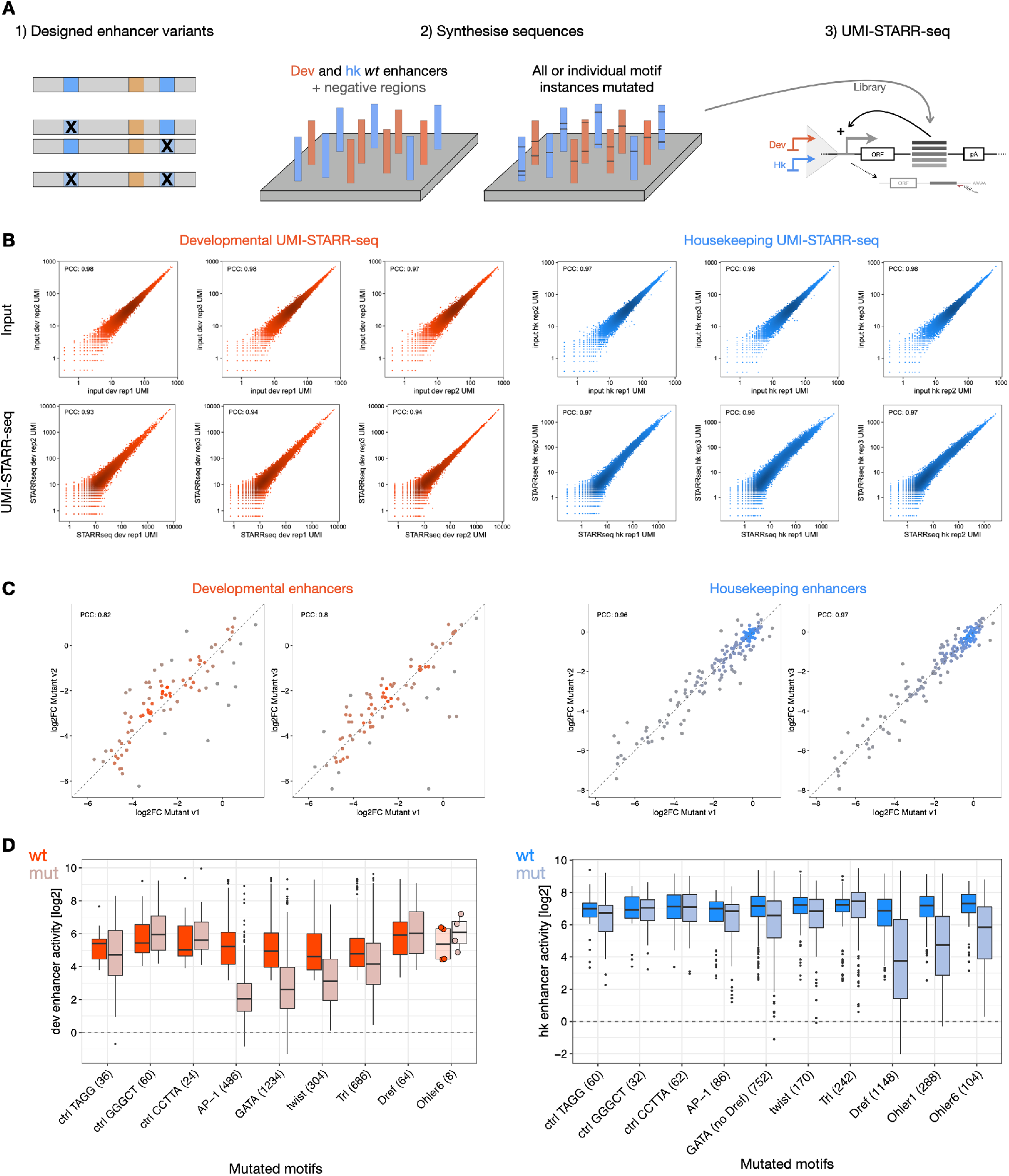
Large-scale systematic TF motif mutagenesis. **A)** Overview of the (1) design, (2) synthesis and (3) UMI-STARR-seq screen of the mutagenesis oligo library. UMI-STARR-seq was performed with a developmental (red) and a housekeeping (blue) promoter in *D. melanogaster* S2 cells. **B)** Pairwise comparisons of input (top) and UMI-STARR-seq (bottom) signal between three independent biological replicates across all oligos included in the library with a developmental (left) or housekeeping (right) promoter. Axes show counts per million in logarithmic scale. The PCC is denoted for each comparison. **C)** Motif requirements are independent of motif mutant variants. Pairwise comparisons of log2 fold-change (log2 FC) to wildtype activity between the three motif-mutant shuffled versions across developmental (left) and housekeeping (right) enhancers. The PCC is denoted for each comparison. **D)** Activity (log2) of wildtype and motif-mutant developmental (left) and housekeeping (right) enhancers that were used to derived the log2 fold-changes from Fig 2C. Number of enhancers mutated for each motif type are shown. The box plots mark the median, upper and lower quartiles and 1.5× interquartile range (whiskers); outliers are shown individually.

**Supplementary Figure 4.**
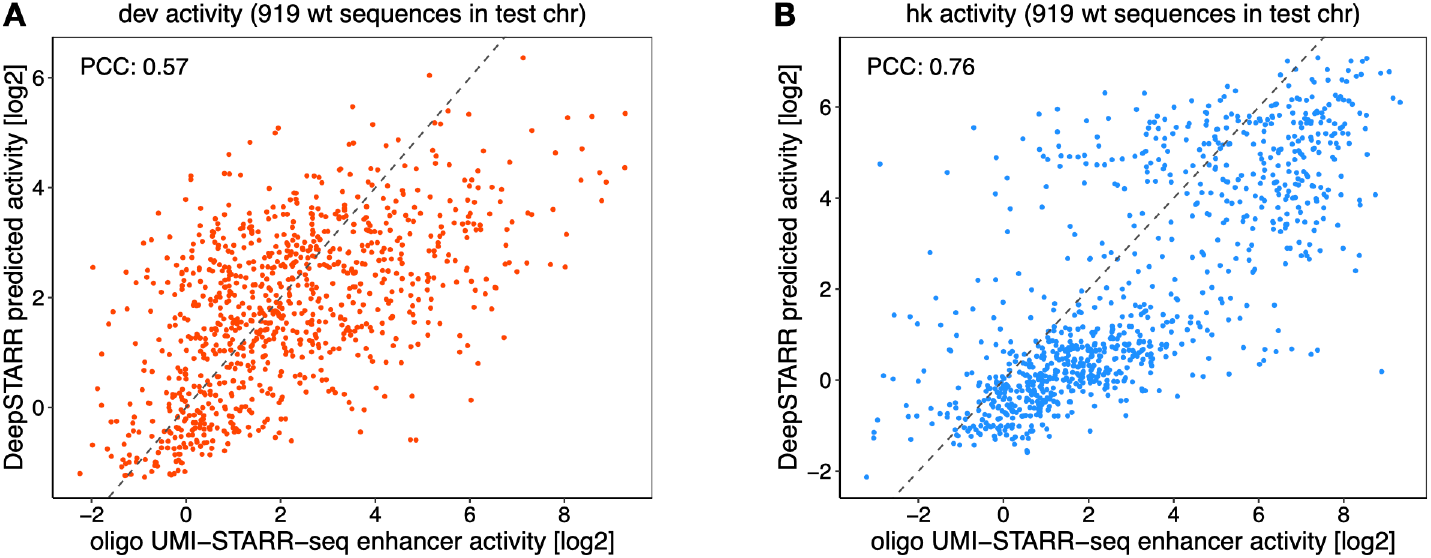
DeepSTARR predicts enhancer activity of wildtype sequences in oligo UMI-STARR-seq. Scatter plots of predicted vs. observed developmental **(A)** and housekeeping **(B)** enhancer activity signal across wildtype sequences from the test set chromosome. The PCC is denoted for each comparison.

**Supplementary Figure 5.**
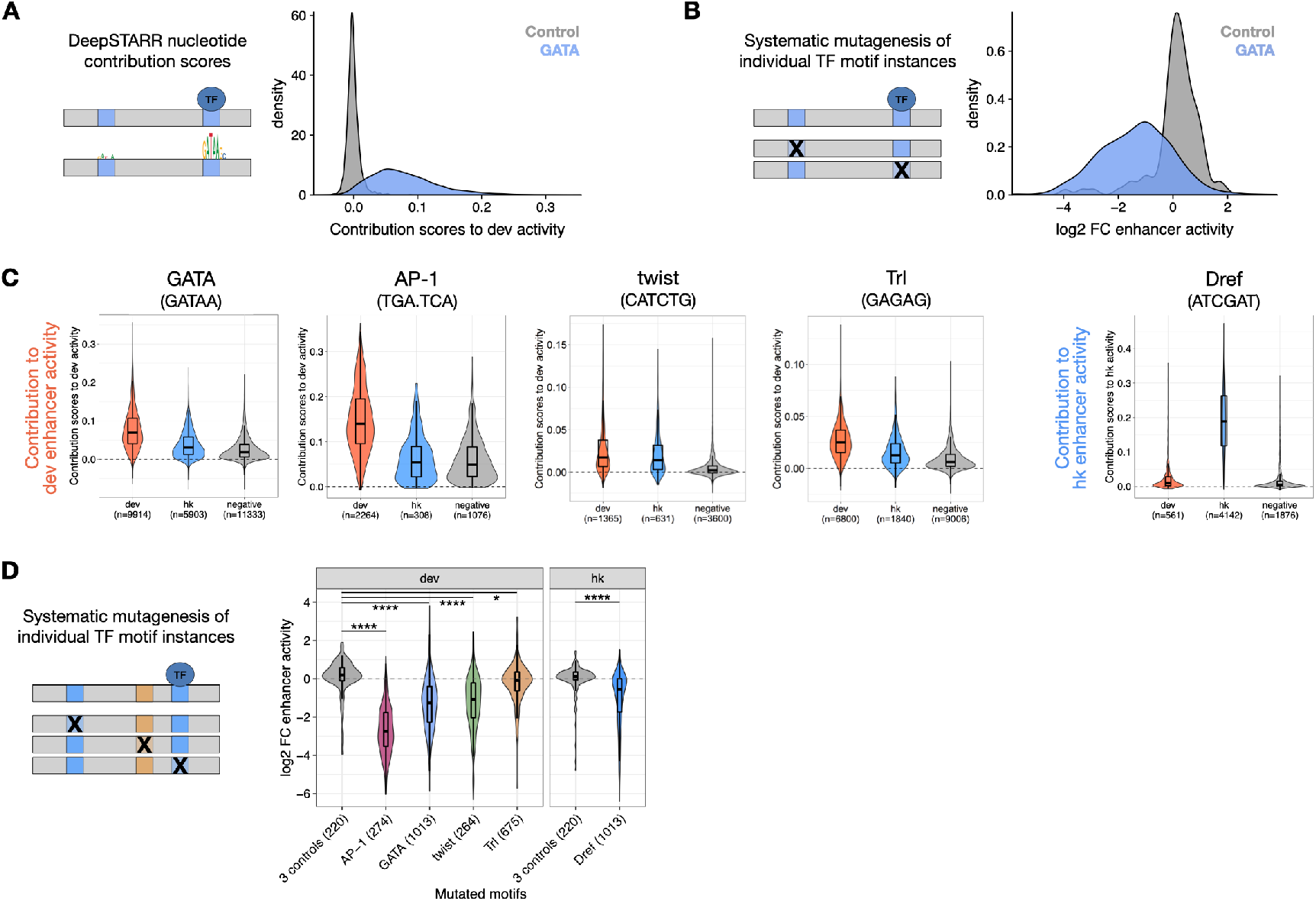
Instances of the same TF motif do not have equivalent contribution to enhancer activity. **A)** DeepSTARR predicts that instances of the same TF motif do not have equivalent contribution. Density distributions of the DeepSTARR predicted contribution scores (average over all its nucleotides) of GATA (blue) or GGGCT (as control; grey) instances in developmental enhancers. **B)** Systematic mutagenesis of individual TF motif instances validates motif non-equivalency. Density distributions of the experimentally derived (oligo UMI-STARR-seq) log2 FC in enhancer activity after mutation of GATA (blue) or control (grey) individual instances in developmental enhancers. **C)** DeepSTARR predicts that instances of the same TF motif are not equivalent. Distributions of the DeepSTARR predicted contribution scores (average over all its nucleotides) of instances of different TF motif types across developmental enhancers (red), housekeeping enhancers (blue) and negative genomic regions (grey). Number of instances for each motif type are shown. The box plots mark the median, upper and lower quartiles and 1.5× interquartile range (whiskers). **D)** Motif mutagenesis validates motif non-equivalency. Distributions of the experimentally derived (oligo UMI-STARR-seq) log2 FC in enhancer activity after mutation of individual instances of different TF motif types or control motifs in developmental or housekeeping enhancers. Note that the core sequence of different instances of the same motif type are identical, despite the different log2 FC. Number of instances for each motif type are shown. The Fligner-Killeen test of homogeneity of variances was used to compare the distributions of each TF motif type with the one from control motifs: **** p-value < 0.0001 and * < 0.05. Box plots as in (C).

**Supplementary Figure 6.**
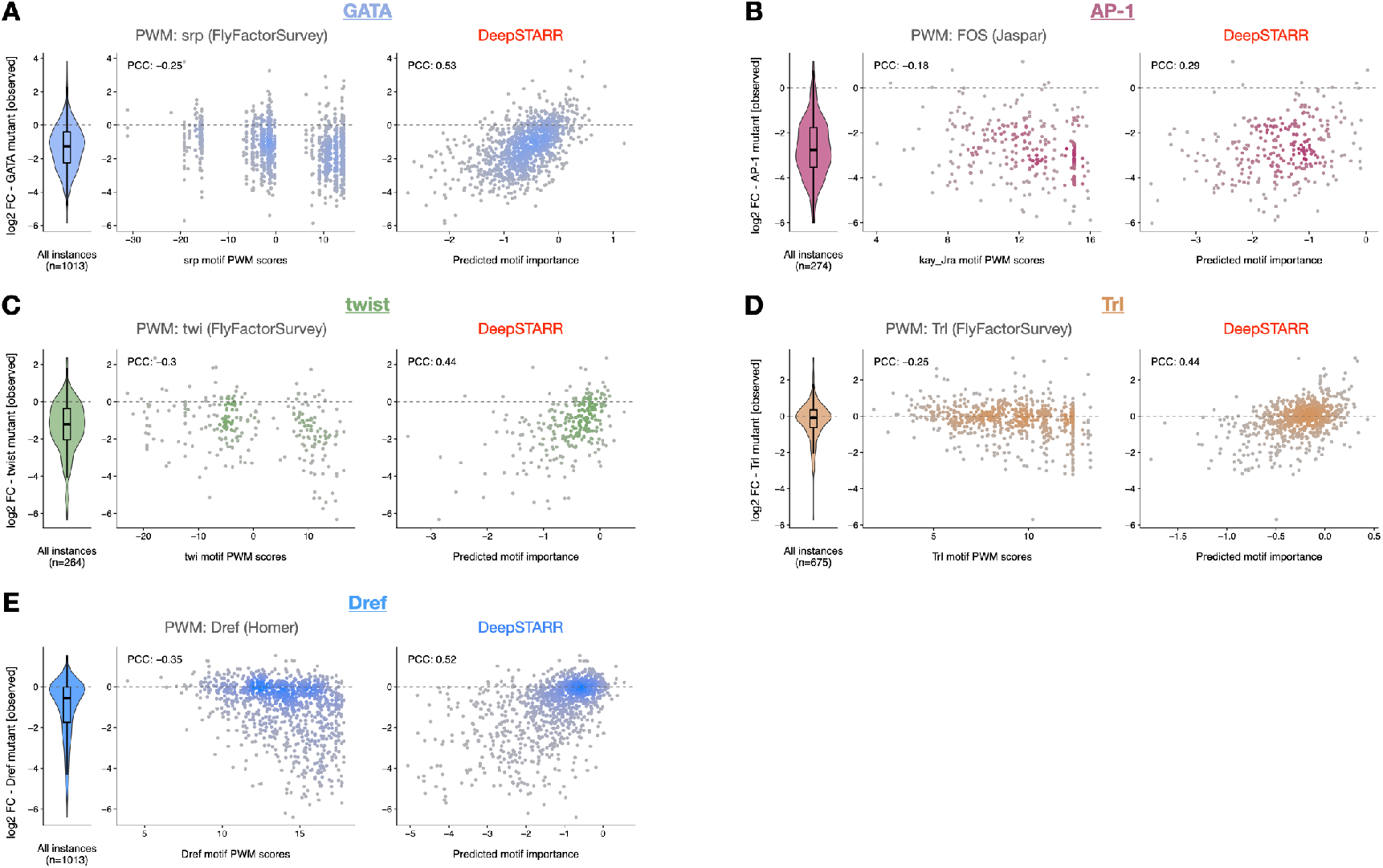
Prediction of motif contribution by PWM scores or DeepSTARR. Distribution of experimentally measured fold-change (log2 FC) enhancer activity after mutating individual motif instances of the GATA **(A)**, AP-1 **(B)**, twist **(C)**, Trl **(D)** and Dref **(E)** motifs (violin plots), compared with the respective TF motif PWM scores and the log2 FC predicted by DeepSTARR. The PCC is denoted for each comparison. The box plots mark the median, upper and lower quartiles and 1.5× interquartile range (whiskers).

**Supplementary Figure 7.**
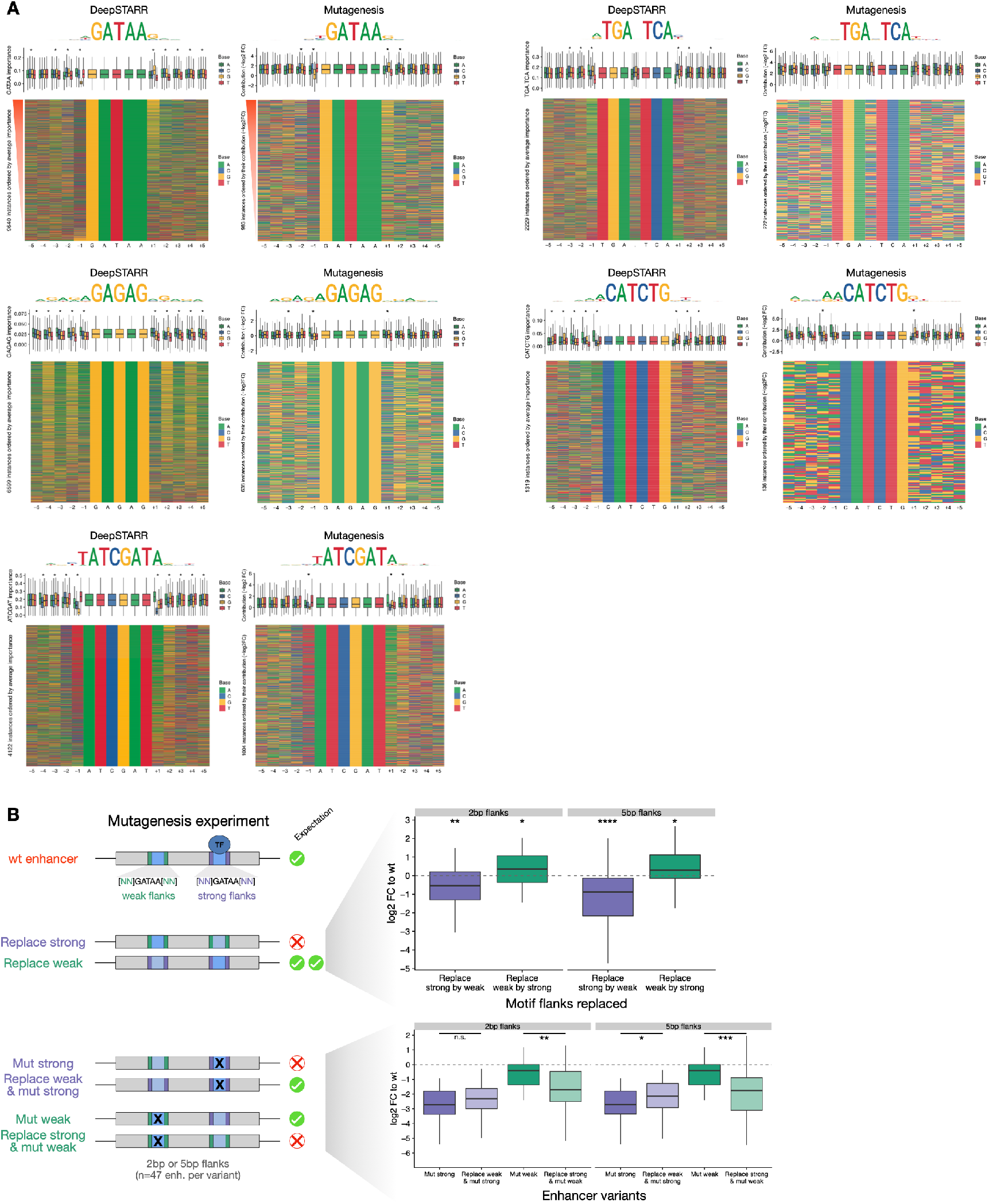
Contribution of TF motifs depend on their flanks. **A)** Motif contribution correlates with flanking base-pairs. Heatmaps: Flanking nucleotides of instances of different TF motif types across developmental (GATA: GATAA, AP-1: TGA.TCA, Trl: GAGAG, twist: CATCTG) or housekeeping (Dref: ATCGAT) enhancers sorted by their DeepSTARR predicted contribution (left) or the experimentally derived (oligo UMI-STARR-seq) log2 fold-change in enhancer activity after mutation (right; minus log2 fold-change, -log2 FC). Box plots: Importance of motif instances according to the different bases at each flanking position. * marks positions with significant differences between the four nucleotides (FDR-corrected Welch One-Way ANOVA test p-value < 0.01). The box plots mark the median, upper and lower quartiles and 1.5× interquartile range (whiskers). Number of instances for each motif type are shown. Top: logos of the top 90^th^ percentile motif instances for each sorting method. **B)** GATA flanking nucleotides are sufficient to switch motif contribution. 47 developmental enhancers containing both one strong (purple) and one weak (green) GATA instance (≥ 2-fold difference between instances) were selected. Top: log2 FC enhancer activity to wildtype for sequences where the 2 or 5 bp flanks of strong instances were replaced by the ones of weak instances (purple) and vice versa (green). Bottom: log2 FC enhancer activity to wildtype of mutating the strong instance (purple) compared to mutating this instance and additionally replacing the 2 or 5 bp flanks of the weak instance by the flanks of the strong instance (light purple). Log2 FC of mutating the weak instance (green) compared to mutating this instance and additionally replacing the 2 or 5 bp flanks of the strong instance by the flanks of the weak instance (light green). **** p-value < 0.0001, *** < 0.001, ** < 0.01, * < 0.05 (Wilcoxon signed rank test). Box plots as in (A).

**Supplementary Figure 8.**
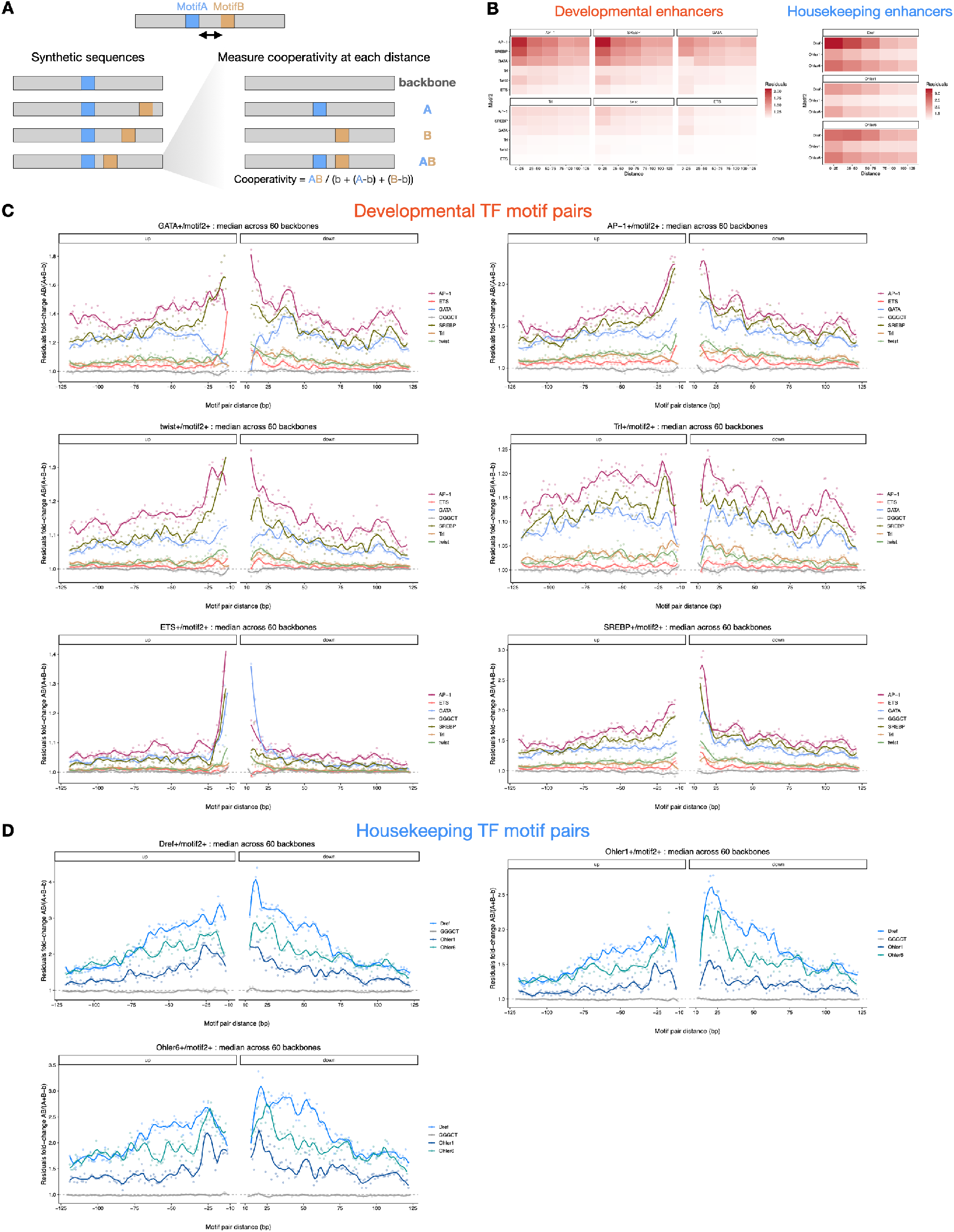
Interpretation of DeepSTARR reveals TF motif distance preferences. **A)** *In silico* characterization of TF motif distance preferences. *MotifA* was embedded in the center of 60 synthetic random DNA sequences and *MotifB* at a range of distances from *MotifA*, both up- and downstream. Both the average developmental and housekeeping enhancer activity is predicted by DeepSTARR. The cooperativity (residuals fold-change) between *MotifA* and *MotifB* as a function of distance is quantified as the activity of *MotifA+B* divided by the sum of the marginal effects of *MotifA* and *MotifB* (*MotifA* + *MotifB –* backbone (b)). **B)** Heatmaps showing the pairwise cooperativity (residuals) between different TF motif types in developmental (left) or housekeeping (right) enhancers. **C-D)** Cooperativity between motif pairs at different distances in (C) developmental and (D) housekeeping enhancers. Points and smooth lines show the median cooperativity across all 60 backbones for each motif pair distance up- and downstream. The *MotifA* in the center is mentioned in each plot’s title and tested with all *MotifB* motifs (different colours). GGGCT motif was used as control (grey). Dashed line at 1 represents no interaction.

**Supplementary Figure 9.**
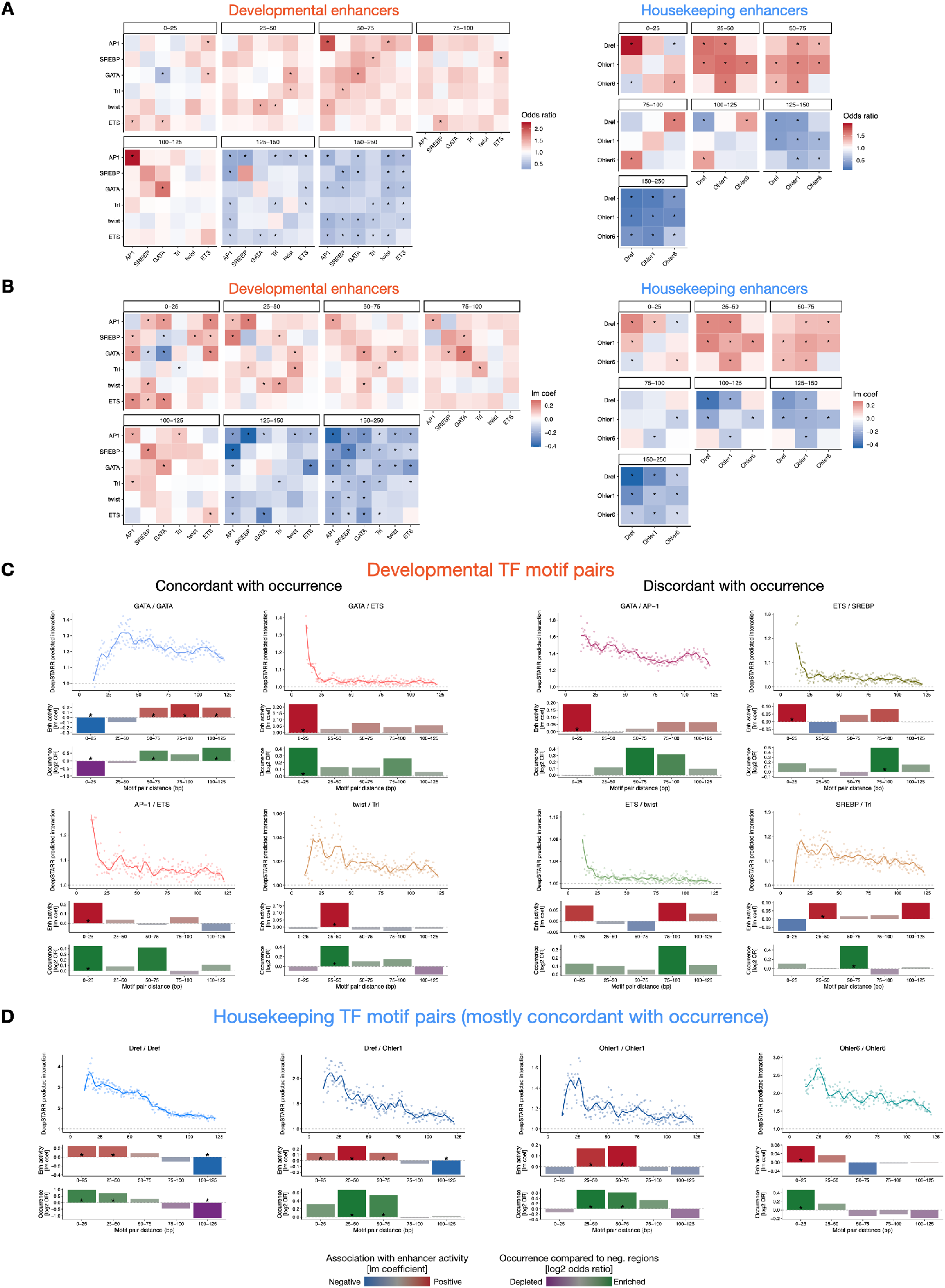
Motifs are not often at optimal distances in developmental enhancers, but enhancer activity follows optimal spacing rules. **A)** Occurrence of motif pairs at different distances in genomic enhancers. Heatmaps showing the enrichment (Fisher’s odds ratio) of motif pairs at different distance bins in developmental (left) or housekeeping (right) enhancers. * represents significant enrichment or depletions (FDR-corrected p-value < 0.05). **B)** Validation of optimal spacing rules for enhancer activity. Heatmaps showing the association between enhancer activity and the presence of motif pairs at different distance bins in developmental (left) or housekeeping (right) enhancers using a multiple linear regression. The multiple linear regression included, as independent variables, the number of instances for the different developmental or housekeeping TF motif types. Linear model coefficients are shown and * represents significant positive or negative associations (FDR-corrected p-value < 0.05). **C-D)** Top: Same as in Fig S8C,D (but with up- and downstream distances combined) per (C) developmental or (D) housekeeping motif pair. Middle: Association between enhancer activity and the distance at which the motif pair is found. Coefficient (y-axis) and p-value from a multiple linear regression including, as independent variables, the number of instances for the different developmental or housekeeping TF motif types. Bottom: Odds ratio (log2) by which the two motifs are found within a specified distance from each other in enhancers compared with negative genomic regions. Color legend is shown. Example motif pairs where optimal spacing preferences are concordant or discordant with their occurrence in enhancers are shown. * FDR-corrected Fisher’s Exact test p-value < 0.05.

**Supplementary Figure 10.**
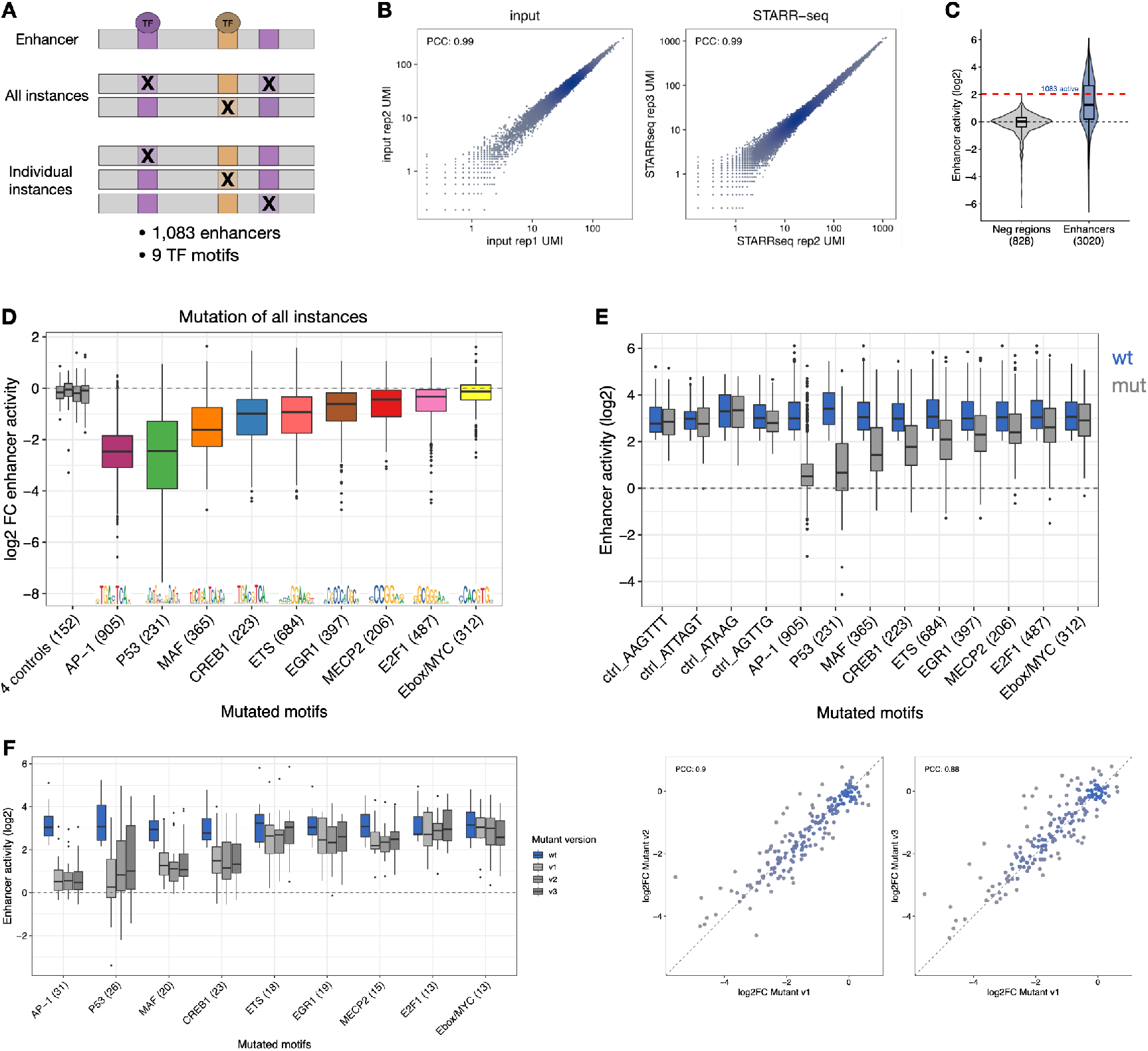
Systematic TF motif mutagenesis in human HCT116 enhancers. **A)** Systematic TF motif mutagenesis in human HCT116 enhancers. We selected 1,083 strong human enhancers and 9 TF motif types and mutated all instances of the same motif simultaneously or each instance individually. The activity of the wildtype and mutant sequences were measured through UMI-STARR-seq. **B)** Pairwise comparisons of input and STARR-seq signal between two independent biological replicates across all oligos included in the human oligo library. Axes are in logarithmic scale. The PCC is denoted for each comparison. **C)** Identification of 1,083 active short human enhancers. Distribution of log2 enhancer activity for oligos selected from negative regions (grey) or enhancer sequences (blue). 1,083 active short human enhancers (log2 wildtype activity in oligo UMI-STARR-seq >= 2.03, the strongest negative region, red dashed line; see Methods) were selected for subsequent motif mutation analyses. The box plots mark the median, upper and lower quartiles and 1.5× interquartile range (whiskers). **D)** TF motif requirements of human HCT116 enhancers. Log2 FC enhancer activity for hundreds of human enhancers after mutating all instances of four control (grey) and nine candidate human TF motifs. Number of enhancers mutated for each motif type and respective motif PWM logos are shown. Box plots as in (C); but outliers are shown individually. **E)** Activity (log2) of wildtype and motif-mutant enhancer sequences that were used to derived the log2 fold-changes from Fig S10D. Number of enhancers mutated is shown. Box plots as in (C); but outliers are shown individually. **F)** Motif requirements are independent of motif mutant variants. Left: Distribution of enhancer activity for wildtype or motif-mutant enhancer sequences for the different TF motifs. The activity of sequences where the motifs were mutated to different motif shuffled versions is shown. Number of enhancers mutated for each motif type are shown. Box plots as in (C); but outliers are shown individually. Right: Pairwise comparisons of log2 FC to wildtype activity between the three motif-mutant shuffled versions across all enhancers. The PCC is denoted for each comparison.

**Supplementary Figure 11.**
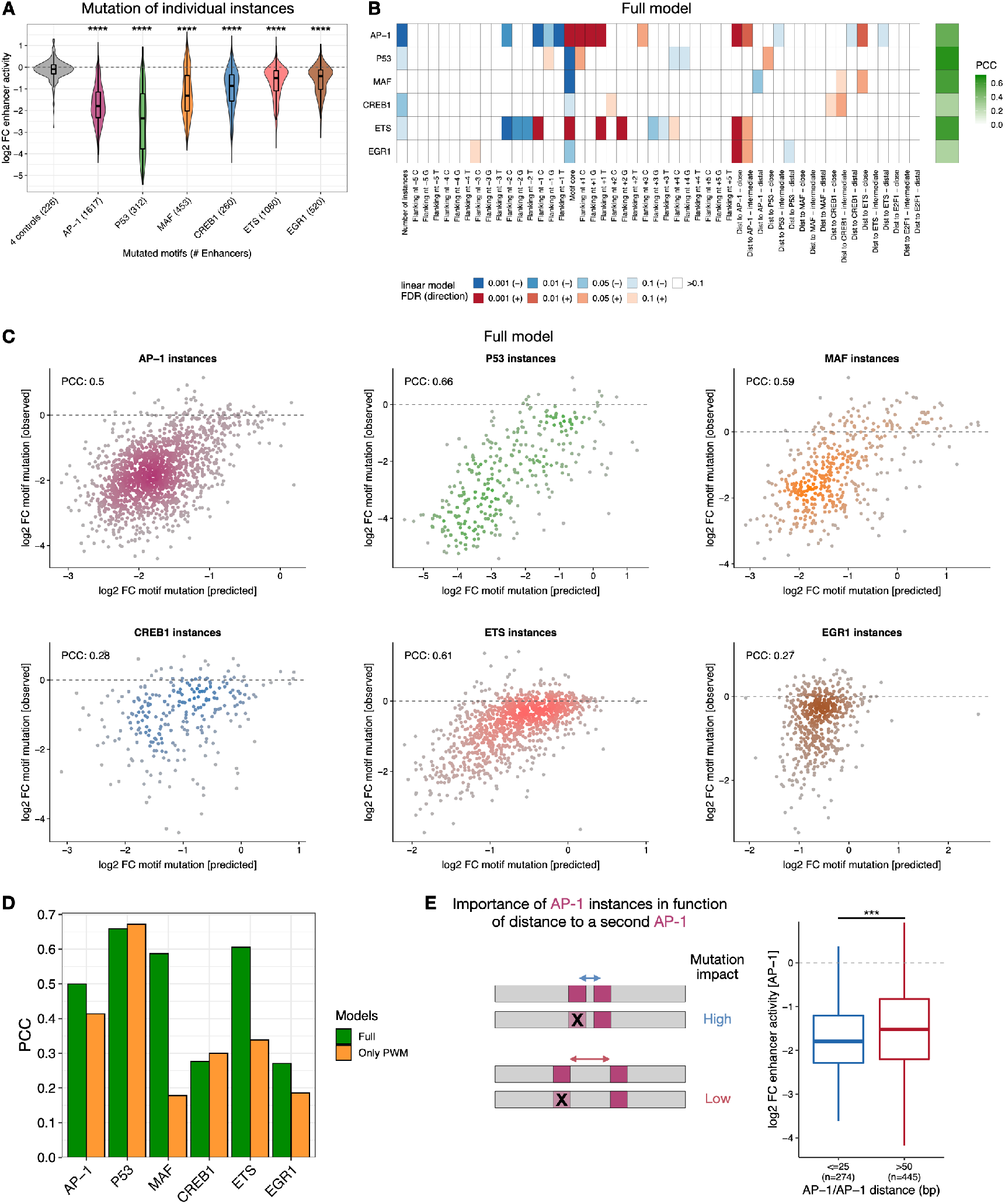
Motif syntax rules dictate the contribution of motif instances. **A)** TF motif non-equivalence is widespread in human enhancers. Distributions of the log2 FC in enhancer activity after mutation of individual instances of different TF motif types or control motifs. Number of instances for each motif type are shown. The Fligner-Killeen test of homogeneity of variances was used to compare the distributions of each TF motif type with the one from control motifs: **** p-value < 0.0001. The box plots mark the median, upper and lower quartiles and 1.5× interquartile range (whiskers). **B)** Motif syntax rules dictate the contribution of TF motif instances in human enhancers. For each TF motif type (rows), we built a linear model containing the number of instances, the motif core (defined as the nucleotides included in each TF motif PWM model) and flanking nucleotides (5 bp on each side), and the distance to all other TF motifs (close: < 25 bp; intermediate: ≥ 25 bp and ≤ 50 bp; distal: >50 bp) to predict the contribution of its individual instances (mutation log2 FC, from Fig S11A) across all enhancers. Heatmap shows the contribution of each feature (columns) for each model, colored by the direction (positive: red, negative: blue) and FDR-corrected p-value. The PCC between predicted and observed motif contribution is shown with the green color scale. **C)** Scatter plots comparing the measured contribution of individual instances of each TF motif type (log2 FC in enhancer activity after mutation) with the one predicted by the models from (B). The PCC is denoted for each comparison. **D)** Models taking into the motif syntax features predict better the contribution of motif instances than solely the PWM scores. Bar-plots comparing the PCC from the full models (from (B); green) and the same just using existing PWM scores (orange). **E)** Motif mutagenesis validates that AP-1 instances close to a second AP-1 instance are more important. Left: expected mutational impact when mutating AP-1 instances depending on the distance to other AP-1 motifs. Right: enhancer activity changes (log2 FC) after mutating AP-1 instances at optimal close (< 25 bp) or suboptimal longer (> 50 bp) distance to a second instance. Number of instances are shown. *** p-value < 0.001 (Wilcoxon rank-sum test). Box plots as in (A).

**Supplementary Figure 12.**
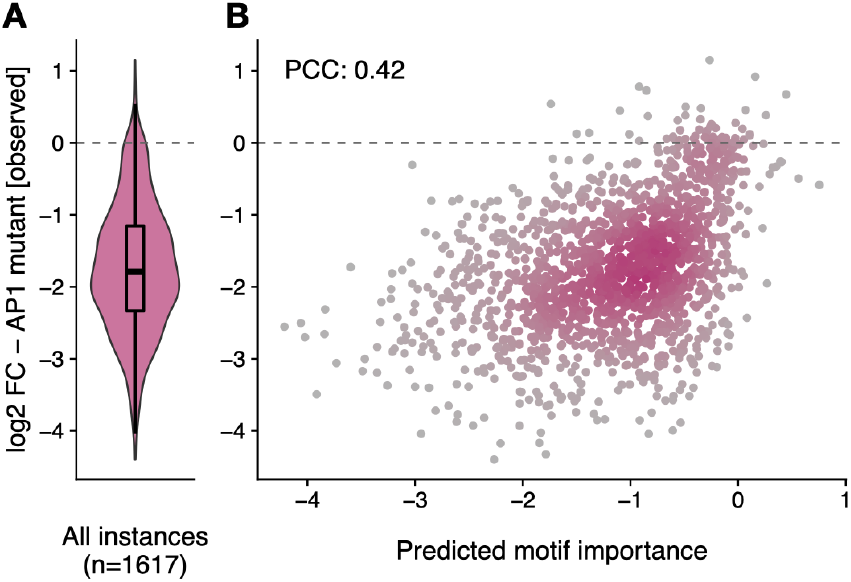
DeepSTARR predicts the contribution of AP-1 instances in human enhancers. Distribution of experimentally measured log2 fold-change (log2 FC) enhancer activity after mutating 1,617 different AP-1 instances across HCT116 enhancers **(A)**, compared with the log2 FC predicted by DeepSTARR **(B)**. The PCC is denoted. The box plots mark the median, upper and lower quartiles and 1.5× interquartile range (whiskers).

